# Depletion of CD206^+^ Tumour Macrophages via a Peptide-Targeted Star-Shaped Polyglutamate Inhibits Tumourigenesis and Metastatic Dissemination in Breast Cancer Models

**DOI:** 10.1101/2021.12.29.474487

**Authors:** Anni Lepland, Alessio Malfanti, Uku Haljasorg, Eliana K. Asciutto, Monica Pickholz, Mauro Bringas, Snežana Đorđević, Liis Salumäe, Pärt Peterson, Tambet Teesalu, María J. Vicent, Pablo Scodeller

**Author notes:** equal contribution (Anni Lepland, Alessio Malfanti). equal contribution (Tambet Teesalu, María J. Vicent, Pablo Scodeller).

## Abstract

Although many studies have explored the depletion of tumour-associated macrophages (TAMs) as a therapeutic strategy for solid tumours, currently available compounds suffer from poor efficacy and dose-limiting side effects. Here, we developed a novel TAM-depleting agent (“OximUNO”) that specifically targets CD206^+^ TAMs and demonstrated efficacy in triple negative breast cancer (TNBC) mouse models. OximUNO comprises a star-shaped polyglutamate (St-PGA) decorated with the CD206-targeting peptide mUNO that carries the chemotherapeutic drug doxorubicin (DOX). In TNBC models, a fluorescently labelled mUNO-decorated St-PGA homed to CD206^+^ TAMs within primary lesions and metastases. OximUNO exhibited no acute liver or kidney toxicity in vivo. Treatment with OximUNO reduced the progression of primary tumour lesions and pulmonary metastases, significantly diminished the number of CD206^+^ TAMs and increased the CD8/FOXP3 expression ratio (demonstrating immunostimulation). Our findings suggest the potential benefit of OximUNO as a TAM-depleting agent for TNBC treatment. Importantly, our studies also represent the first report of a peptide-targeted St-PGA as a targeted therapeutic nanoconjugate.

## INTRODUCTION

Triple negative breast cancer (TNBC), defined by the lack of the expression of the oestrogen receptor (ER), progesterone receptor (PR), and human epidermal growth factor receptor 2 (HER2)^1,2^, represents an aggressive breast cancer subtype with poor prognosis^3^ that comprises up to 20% of all breast cancer cases^3,4^. Interfering with immune checkpoints signalling (e.g. through the modulation of programmed cell death 1 (PD-1) and its ligand (PD-L1)) represents an alternative treatment strategy for several cancers and is currently being employed in combination with chemotherapy as a neoadjuvant or adjuvant treatment^5–8^. The U.S. Food and Drug Administration (FDA) recently granted accelerated approval for a combination of a PD-L1 blocking antibody (atezolizumab, Tecentriq®) and nab-paclitaxel (Abraxane®)^9^ as a first-line treatment for unresectable locally advanced or metastatic TNBC^10^. While promising clinical results have resulted, this combinatorial treatment approach suffers from significant obstacles, including the problematic identification and heterogeneity of PD-L1 expression in patients^11^, the limited applicability to PD-L1 positive TNBC patients (only 20–42% of cases)^12,13^, and the induction of severe side effects (e.g., neutropenia, peripheral neuropathy, and colitis)^10,14,15^. Other immune checkpoint inhibitors (ICIs), including the cytotoxic T lymphocyte-associated antigen 4 (CTLA-4) blockers ipilimumab and tremelimumab, are currently under evaluation for TNBC treatment in combination with other drugs (clinical trial identifiers: NCT03606967, NCT02983045); however, anti-CTLA-4 treatments induce severe side effects such as endocrinopathies, myopathy, enterocolitis, and hepatitis^16–19^, which narrow their use. Overall, the limited success of alternative treatment options for TNBC has maintained chemotherapy as the standard of care for most patients^20^.

The anthracycline drug doxorubicin (DOX), which presents high off-target effects such as cardiotoxicity^21,22^, represents a frequently employed chemotherapeutic for TNBC; however, disease relapse and metastatic development have also been associated with DOX treatment^23^. M2 (anti-inflammatory)-polarised tumour-associated macrophages (TAMs)^24^ found within both primary and metastatic tumour lesions mediate both events^25^; furthermore, TAMs represent the main executioners of tumour progression, immunosuppression and invasion^24–29^, and their presence correlates with inadequate therapeutic response and poor prognosis^25^. Recent efforts have focused on eliminating TAMs, and several ongoing clinical trials are currently evaluating TAM depletion in combination with treatments such as ICIs^30^. The current clinical-stage gold standard for TAM depletion relies on agents that block colony stimulating factor 1 (CSF1) or its receptor CSF1R, such as the small molecule CSF1R inhibitor PLX3397^31^; however, microglia also expresses CSF1R^32^, the inhibition of CSF1R with PLX5622 impacts M1 macrophages^33^, and PLX3397 treatment causes oedema^34^. Clinical data suggests that anti-CSF1R antibodies induce a modest effect^35,36^ and cause severe side effects that include haematological toxicities^35^ and hepatotoxicity by targeting Kupfer cells^35,36^. Overall, these findings highlight the overwhelming need for new TAM-depletion strategies.

Notably, both perivascular TAMs associated with disease relapse and therapeutic resistance^24^ and metastasis-associated macrophages^37^ express the mannose receptor (CD206/MRC1). Perivascular TAMs employ CD206 to navigate the surrounding collagen-dense stroma^38^, which favours tumour progression^39,40^.

For the first time, we report the effects of depleting the CD206^+^ subpopulation of TAMs in metastatic TNBC mouse models through the use of a targeting agent (the mUNO peptide) for a CD206 site different from the mannose-binding site^41–44^. Previous studies have employed mannose to target CD206; however, mannose has other receptors besides CD206^45,46^.

We decorated a three-arm branched biodegradable multivalent polyanion with a defined negative charge and nanometre-size hydrodynamic radius (star-shaped polyglutamate or St-PGA) with mUNO peptide to function as a targeted delivery platform for a chemotherapeutic agent (DOX) conjugated through a bioresponsive linker. St-PGA-DOX-mUNO (referred to as OximUNO) efficiently depleted CD206^+^ TAMs, relieved immunosuppression in the tumour microenvironment (TME) and limited metastasis/tumour growth, thereby supporting OximUNO as an alternative TAM depletion strategy.

Most importantly, this study represents the first described combination of two reported technologies – the St-PGA nanocarrier and the mUNO targeting peptide. Overall, this OximUNO proof-of-concept demonstrates the potential of the peptide-targeted St-PGA nanosystem. Our studies lay a foundation for future work using this nanosystem to target other receptors efficiently by changing the targeting peptide.

## RESULTS

### Design and structural modelling of St-PGA-OG-mUNO

To characterise and explore the function of OximUNO, we first developed an mUNO-targeted St-PGA labelled with the Oregon Green (OG) fluorescent dye (referred to as St-PGA-OG-mUNO) (Fig. 1A, Scheme S1). We conjugated OG to St-PGA using an amide linker to allow in vitro or in vivo tracking and coupled mUNO through a disulphide bond formed between the free cysteine of mUNO and a pyridyldithiol linker on St-PGA. We previously demonstrated that mUNO conjugated to polymeric nanostructures through the cysteine thiol group preserves CD206 binding^42^. To evaluate the structure and dye loading, we analysed St-PGA-OG-mUNO and St-PGA-OG using nuclear magnetic resonance (NMR) and UV-Vis analyses (Fig. S1).

**Figure 1.**
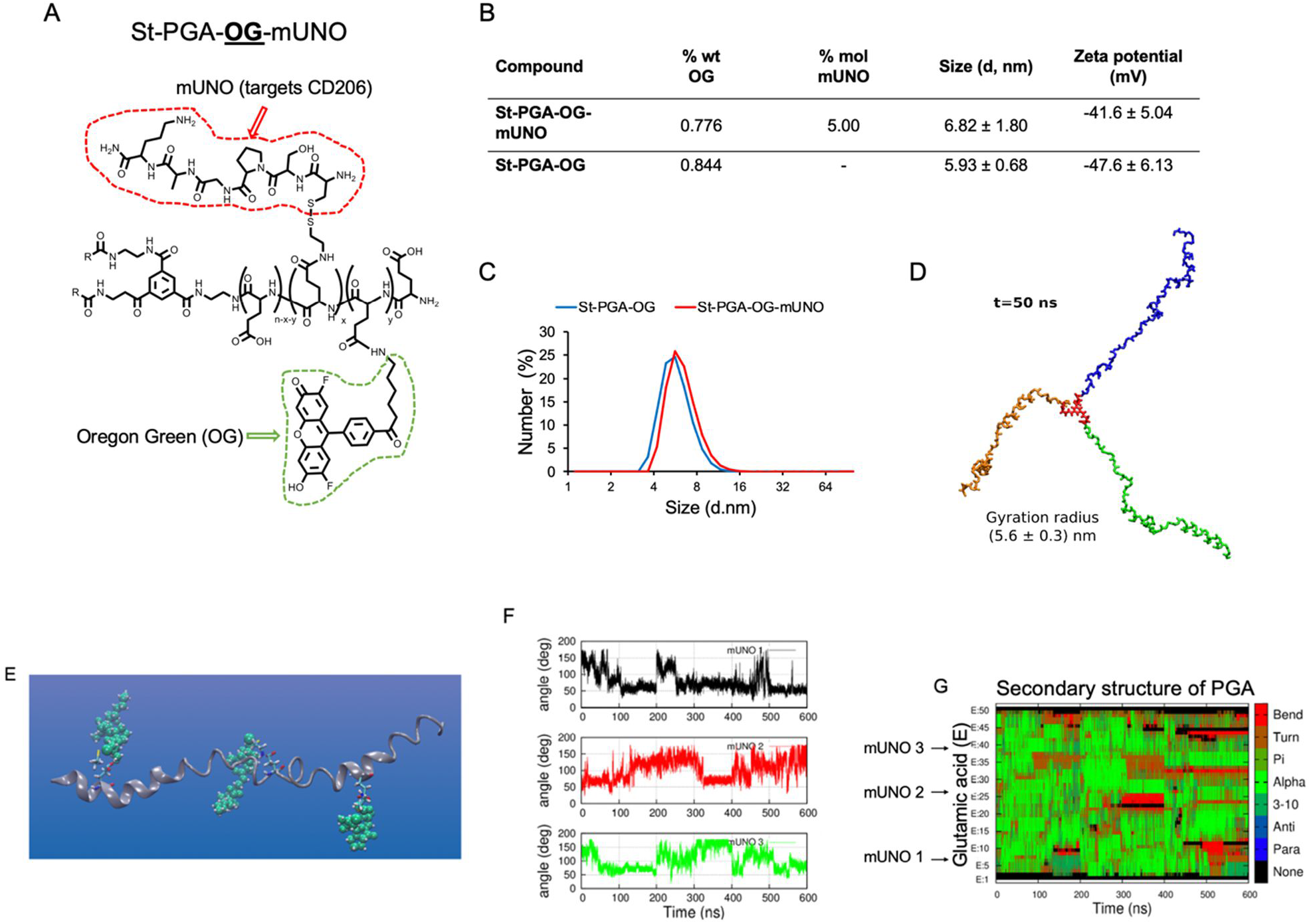
Design and analysis of mUNO-targeted St-PGA. (**A**) Representative structure of St-PGA decorated with mUNO peptides (red) and OG (green). (**B**) Table detailing OG loading, mUNO loading, size (as measured by DLS), and charge (as measured by Zeta potential analysis). (**C**) DLS graph demonstrating uniform size for both St-PGA-OG-mUNO and St-PGA-OG. (**D**) A snapshot of modelled St-PGA structure in water and Na^+^ counterions at the last stage of the simulation (50 ns), displaying the three arms in different colours for visual clarity. The average gyration radius was 5.6 ± 0.3nm, t shows time in ns. (**E**) Representative MD snapshot of a single St-PGA-mUNO branch containing three equidistant mUNO peptides. Green spheres represent mUNO and a Licorice representation shows the linker. (**F**) mUNO rotation around the PGA chain for each of the three peptides (black, red, and green lines). (**G**) PGA chain secondary structure evolution, where red and brown regions show how mUNO perturbs the chain structure, turning alpha helices into random coils.

Dynamic light scattering (DLS) analysis demonstrated that St-PGA-OG-mUNO and St-PGA-OG displayed similar hydrodynamic diameters of 6.8 and 5.9 nm, respectively (Fig. 1B, C), while both nanoconjugates exhibited highly negative charges (−42 mV and -48 mV, respectively) as shown by Zeta potential analysis (Fig. 1B); an expected result given the glutamic acid nature of the polymer carrier. Analysis of mUNO loading (Fig. 1B) indicated the presence of approximately seven mUNO peptides in St-PGA-mUNO nanoconjugate, which would allow multivalent receptor binding.

We next assessed the structure of unlabelled and untargeted St-PGA in water using molecular dynamics (MD) simulations to access information at the atomic scale. We assumed an initial helical conformation for the three PGA chains. The studied system consisted of a fully hydrated St-PGA and the Na^+^ counterions (∼920,000 atoms) and was built after initial minimisation under vacuum conditions. We simulated 50 ns of the entire St-PGA macromolecule, with Fig. 1D displaying a snapshot corresponding to the last step of the simulation. Averaging the gyration radius over the last 25 ns of the simulation run provided a value of 5.6 ± 0.3 nm, which lies in the same order of magnitude as the results from DLS analysis and suggests a lack of aggregation of both St-PGA-OG-mUNO and St-PGA-OG in PBS. A video simulation (Video S1) suggested that the three PGA chains remain in an extended conformation throughout the simulation and do not show any intra- or inter-molecular interaction, suggesting that the mUNO peptides linked to St-PGA will not interfere with each other.

To investigate if mUNO can engage with the CD206 receptor when grafted onto St-PGA, we modelled the structure and mobility of St-PGA-mUNO using computational analysis. To attain a computationally feasible system, we simulated only single branches of St-PGA-mUNO. We placed three equidistant mUNO peptides on a PGA single branch and fully solvated the system. We observed that three mUNO peptides remained exposed to the solution available for receptor binding (Fig. 1E). The rotation of mUNO around PGA, tracked by the angle formed by a proline aromatic carbon within mUNO (Fig. S2, green sphere), a pyridyldithiol linker nitrogen (Fig. S2, blue sphere), and a glutamic acid aromatic carbon (Fig. S2, light blue sphere) revealed angles between 50° and 180° (Fig. 1F). This value supports the ability of mUNO peptides to interact with their receptor^43^. Comparisons with an undecorated PGA branch demonstrated the minimal alterations of secondary structure dynamics in the presence of mUNO peptides - turning alpha helices (Fig. 1G, green) into random coils (Fig. 1G, brown) at regions where they are placed; however, the PGA chain structure remained mainly helical except in the middle, where a slight kink formed (Fig. 1G).

Altogether, St-PGA-OG-mUNO and St-PGA-OG nanoconjugates possessed similar sizes by DLS, highly negative charges, and, according to simulations, displayed their three arms in an extended open structure. Our simulation analyses demonstrated that mUNO peptides induced a minimal effect on PGA structure and rotated around the PGA chain with considerable freedom. Overall, these findings suggest St-PGA-mUNO as a suitable platform for CD206 targeting.

### St-PGA-OG-mUNO targets CD206^+^ TAMs and displays low hepatic accumulation

We next evaluated the potential of St-PGA-OG-mUNO to target CD206^+^ TAMs in two different TNBC models – an orthotopic TNBC model and an experimental metastasis of TNBC model induced by intravenous (i.v.) injection of 4T1 cells. We administered St-PGA-OG-mUNO or St-PGA-OG intraperitoneally (i.p.), allowed circulation for 6 h, and then analysed tumour homing using confocal fluorescence microscopy. Our previous study provided the rationale for the i.p. administration route, where we demonstrated that the i.p. administered mUNO peptide exhibited a substantially longer half-life than i.v. administered mUNO in the same mice (same strain, sex and age) used in this study^42^.

In the orthotopic TNBC model, we observed a high colocalisation of OG/CD206 (Fig. 2A, yellow signal) with St-PGA-OG-mUNO but a much lower colocalisation of OG/CD206 with non-targeted St-PGA-OG (Fig. 2B) (0.57 and 0.21, respectively (Fig. 2M)). We observed a low level of accumulation of St-PGA-OG-mUNO or St-PGA-OG in the liver (Fig. S3A, B). We employed confocal image acquisition parameters throughout this study to visualise CD206 in the tumour without signal saturation. Given the higher levels of CD206 in the tumour, imaging with associated settings provides low CD206 visualisation in the liver. Using a higher image intensity, we observed the expected CD206 signal in the liver (as expected from Kupfer cells and sinusoid vessels) (Fig. S4A) and a saturated CD206 signal in the tumour (Fig. S4B).

**Figure 2.**
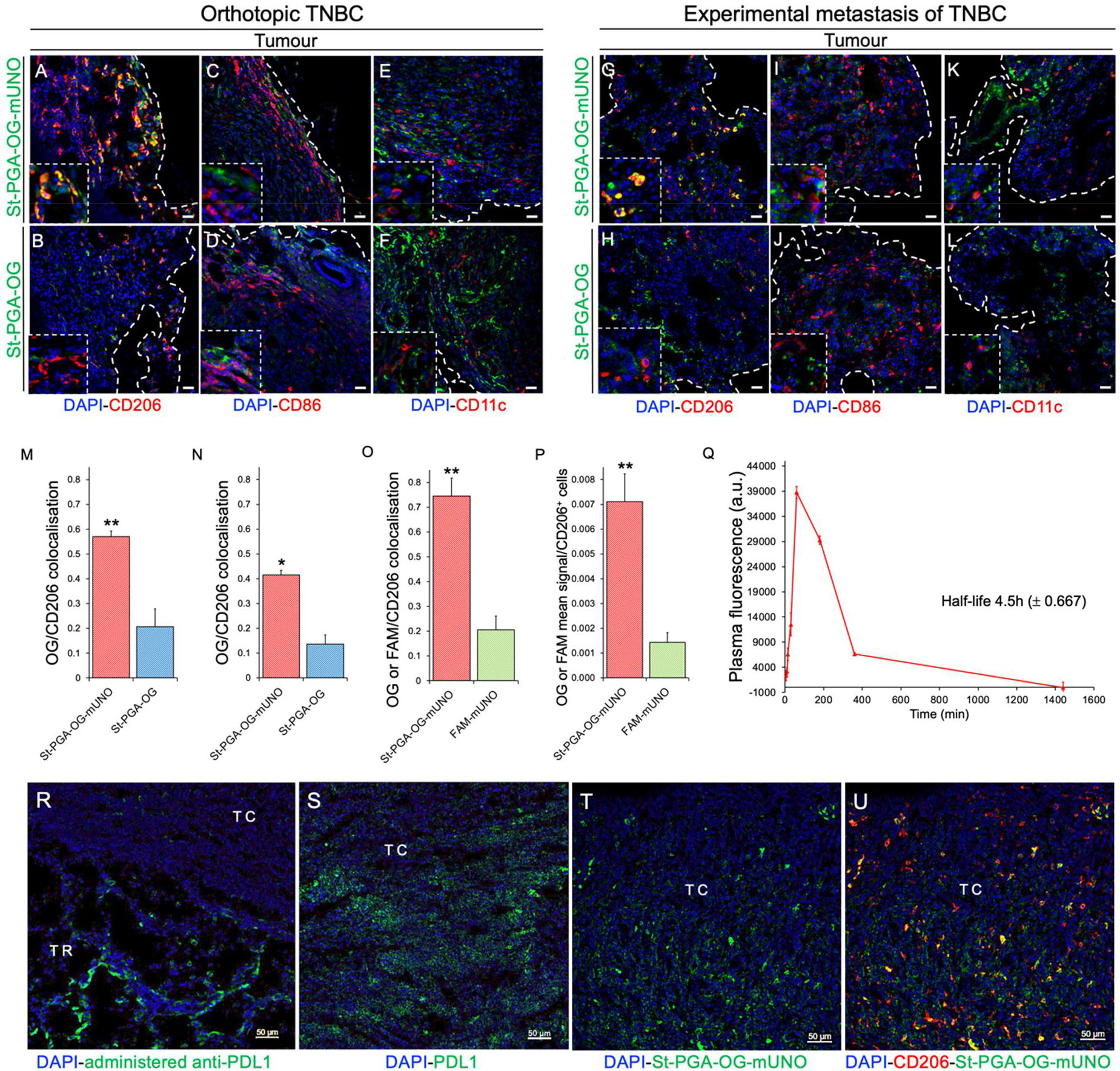
St-PGA-OG-mUNO targets CD206^+^ TAMs in models of orthotopic TNBC and experimental metastasis of TNBC and displays an extended plasma half-life. Homing studies with i.p. administered St-PGA-OG-mUNO (0.41 mg/0.5mL of PBS) or St-PGA-OG (0.35 mg/0.5mL of PBS), after 6 h of circulation. N=3 for orthotopic TNBC and N=2 for experimental metastasis of TNBC. (**A-F**) Homing in orthotopic TNBC. (**A**) St-PGA-OG-mUNO displayed high colocalisation between OG and CD206 (yellow signal), whereas (**B**) St-PGA-OG displayed minimal colocalisation. St-PGA-OG-mUNO and St-PGA-OG did not show any homing to (**C, D**) CD86^+^ cells (M1 macrophages) nor (**E, F**) CD11c^+^ cells (DCs). (**G-L**) Homing study in the experimental metastasis of TNBC. (**G**) St-PGA-OG-mUNO displayed high colocalisation with OG and CD206 (yellow signal), whereas (**H**) St-PGA-OG showed minimal colocalisation. St-PGA-OG-mUNO and St-PGA-OG did not show any homing to (**I, J**) CD86^+^ cells (M1 macrophages) or (**K, L**) CD11c^+^ cells (DCs). Scale bars = 20 µm. (**M**) Graphs depicting the quantification of CD206 and OG colocalisation in the orthotopic TNBC and (**N**) the experimental metastasis of TNBC. (**O**) Quantification of colocalisation analysis for St-PGA-OG-mUNO or FAM-mUNO with CD206 homing after 6 h of circulation, N=2 (30 nmoles in OG and FAM, respectively). Colocalisation was quantified using the Fiji programme and Pearson’s coefficient (for more information see Materials and methods). (**P**) Mean OG/FAM signal per CD206^+^ cell analysed using the ImageJ programme. (**Q**) Plasma fluorescence (in the green channel) of i.p. administered St-PGA-OG-mUNO (dose 15 nmoles in OG) in healthy Balb/c mice (N=3). (**R**) Rat anti-mouse PDL1 was i.v. injected ten days post tumour induction (p.i.) and circulated for 24 h after which time, mice were sacrificed, tumours collected, fixed, and stained for PDL1. **(S)** PDL1 expression was detected in s.c. 4T1 tumours by staining with rat anti-mouse PDL1. (**T, U)** Representative images showing St-PGA-OG-mUNO (0.41mg/0.5mL) i.p. injected ten days post tumour induction and circulated for 6 h after which time, mice were sacrificed, tumours fixed, and then stained for OG and CD206. The OG channel is shown separately in **(T),** and the colocalisation with CD206 for the same image is shown in **(U).** Scale bars = 50 µm. TC: tumour core, TR: tumour rim. Error bars represent the standard error (SE) of the mean.

Immunostaining for endogenous mouse IgG in the tumour and the liver indicated the leaky nature of the tumour vasculature (Fig. S5A) compared to the liver vasculature (Fig. S5B) in the 4T1 model. A leaky tumour vasculature favours the hypothesis that St-PGA-OG-mUNO has a more extended (both in time and space) access to CD206 in the tumour than in the liver. We speculate that the leaky tumour vasculature combined with lower CD206 expression in the liver than the tumour explains the low hepatic accumulation of St-PGA-OG-mUNO. St-PGA-OG-mUNO did not accumulate in the lungs (Fig. S6A) or spleen (Fig. S6B); however, we did observe some accumulation in the sentinel lymph node (SLN) (Fig. S6C) and the kidneys (Fig. S6D). Of note, the observed kidney signal agrees with our prior studies that demonstrated the renal excretion of St-PGA^47^.

Importantly, we did not detect homing to M1 macrophages (CD86^+^) or dendritic cells (CD11c^+^, DCs) with St-PGA-OG-mUNO (Fig. 2C, E) or with St-PGA-OG (Fig. 2D, F). In the experimental metastasis of TNBC model, most of the cellular signal of St-PGA-OG-mUNO associated with CD206^+^ TAMs (Fig. 2G, yellow signal) when compared to St-PGA-OG (Fig. 2H) (OG/CD206 colocalisation 0.42 and 0.14, respectively, Fig. 2N). In this model, we also observed no colocalisation between OG and CD86 (M1 macrophages) (Fig. 2I, J) or OG and CD11c (DCs) (Fig. 2K, L) and the observed hepatic accumulation of St-PGA-OG-mUNO or St-PGA-OG was low (Fig. S7).

One of the rationales behind the design of OximUNO was to increase mUNO targeting through increased avidity and plasma half-life. To evaluate these aspects, we compared the homing of St-PGA-OG-mUNO with a monomeric, carboxyfluorescein-labelled mUNO peptide (FAM-mUNO). We note that even given the different nature of the fluorescent labels (OG on St-PGA-OG-mUNO and fluorescein on FAM-mUNO), we did not use their native fluorescence as a readout; instead, we used an antibody that recognises both FAM and OG; therefore, we do not expect biases from potential differences in FAM and OG emissions.

We discovered that St-PGA-OG-mUNO (Fig. S8A) displayed significantly higher OG/CD206 colocalisation than for FAM/CD206 with FAM-mUNO (Fig. S8B) at 6 h (0.74 vs. 0.21, respectively (Fig. 2O)). Additionally, we found that the OG/FAM mean signal per CD206^+^ cell was five times higher for St-PGA-OG-mUNO than FAM-mUNO (Fig. 2P). These findings suggest that conjugating mUNO to the St-PGA backbone greatly improved receptor binding. Plasma half-life analysis for i.p. administered St-PGA-OG-mUNO revealed a 4.5 h half-life (Fig. 2Q), a value over two times longer than that observed after the i.p. administration of FAM-mUNO in our previous study^42^. Overall, this finding suggests that conjugating peptide onto St-PGA increased the plasma half-life of mUNO peptide, a desirable feature that will improve in vivo ligand targeting.

We next compared tumour homing of St-PGA-OG-mUNO to that of a therapeutic monoclonal antibody by i.v. injecting anti-PDL1 in orthotopic 4T1 tumour-bearing mice and allowing circulation for 24 h. We observed that administered anti-PDL1 accumulated in the tumour rim (Fig. 2R, TR) but not in the tumour core (Fig. 2R, TC) even given expression of the receptor (PDL1) in the tumour core (Fig. 2S, TC). The observed accumulation of St-PGA-OG-mUNO in the tumour core (Fig. 2T, TC) and receptor colocalisation (Fig. 2U), supported the implementation of our platform as an efficient alternative to antibody-based therapies such as anti-PDL1 or antibody-drug conjugates.

Administration of a higher dose of both nanoconjugates (0.82 mg/0.5mL St-PGA-OG-mUNO and 0.7 mg/0.5mL St-PGA-OG) resulted in high CD206^+^ TAM targeting (Fig. S9A) albeit at the expense of higher hepatic accumulation (Fig. S9C). For this reason, we employed lower nanoconjugate doses (0.41 mg/0.5mL and 0.35 mg/0.5mL) for subsequent studies.

Overall, we demonstrated that St-PGA-OG-mUNO, homes to CD206^+^ TAMs in both orthotopic and experimental metastasis of TNBC models with no significant hepatic accumulation. We also established that St-PGA-OG-mUNO does not target M1 macrophages or DCs in the tumour, thereby providing evidence of high specificity for CD206^+^ TAMs.

### OximUNO enhances the in vitro cytotoxicity of DOX on M2 macrophages

St-PGA displays a large surface with multiple sites available for the conjugation of pro-apoptotic or cytotoxic cargoes via bioresponsive polymer-drug linkers^48,49^. To selectively deplete CD206^+^ TAMs, we conjugated an apoptotic chemotherapeutic agent (DOX) to St-PGA-mUNO to form St-PGA-DOX-mUNO (designated “OximUNO”) (Fig. 3A, Scheme S2). We conjugated DOX to St-PGA-mUNO using a hydrazone bond^48^ to allow for site-specific drug release in the acidic milieu of the endosomes/lysosomes^48,50^.

**Figure 3.**
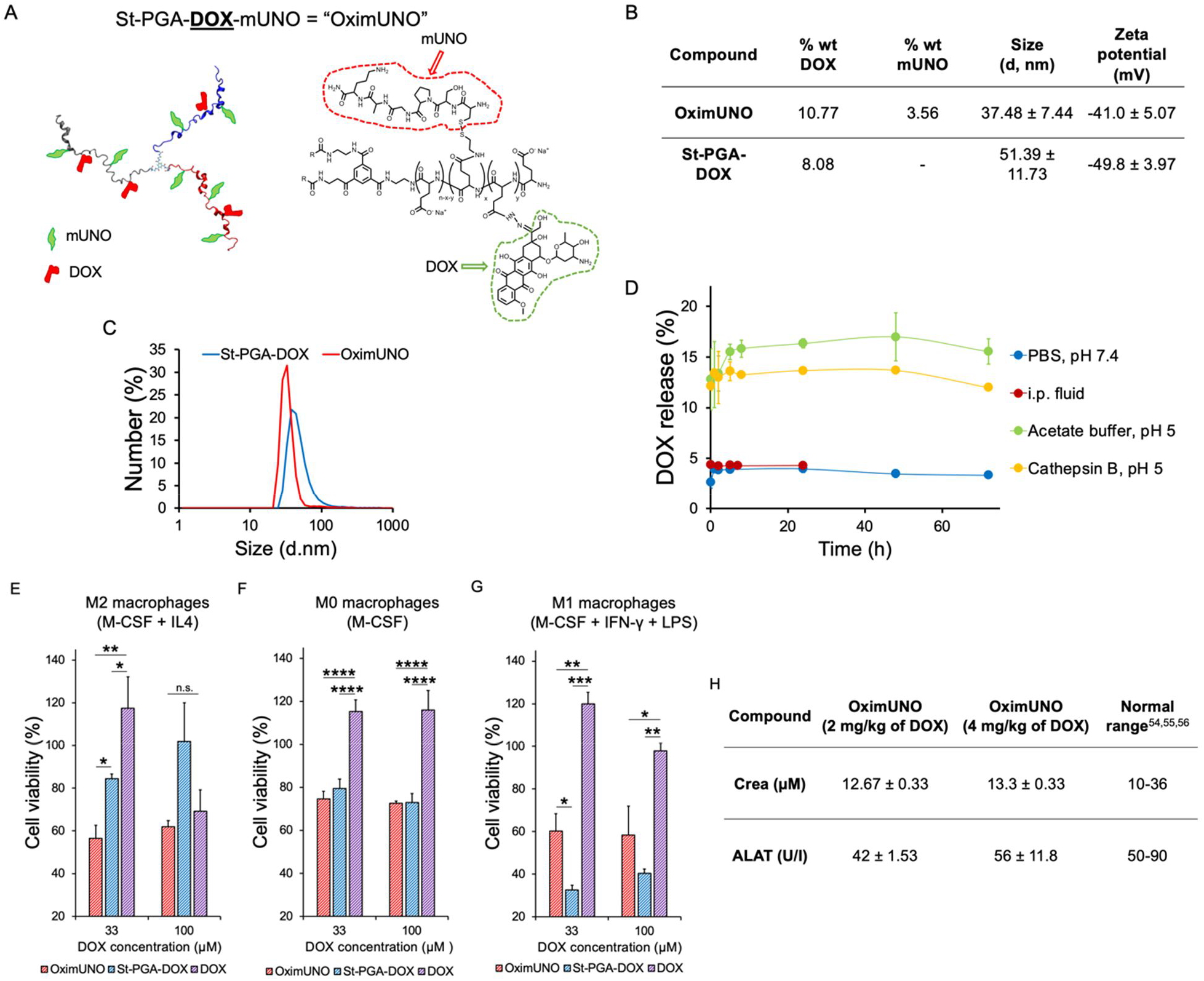
OximUNO enhances the in vitro efficacy of DOX on M2 macrophages. (**A**) Simplified form of OximUNO (left) and molecular structure (right) showing St-PGA decorated with mUNO (red) and DOX (green). (**B**) Table showing DOX loading, mUNO loading, size (as measured by DLS), and charge (as measured by Zeta potential) of both nanoconjugates in PBS. (**C**) A DLS graph for measurements shown in (B), indicating the uniform size of OximUNO and St-PGA-DOX. (**D**) DOX release from OximUNO showing the drug release in PBS, i.p. fluid, acetate buffer or in the presence of cathepsin B. (**E-G**) In vitro cytotoxicity in primary human (**E**) M2, (**F**) M0, or (**G**) M1 macrophages after treatment with OximUNO (red bars), St-PGA-DOX (blue bars), and DOX (purple bars) following a 15 min incubation, washed, cultured for additional 48 h, and then analysed for cell viability as evaluated by MTT assay. (**H**) Hepatic and renal toxicology serum levels of Crea and ALAT 48 h after i.p.-administration of OximUNO (at two different doses in DOX: 2 mg/kg and 4 mg/kg) in healthy Balb/c mice (N=3). Error bars represent the SE of the mean.

To evaluate the effect of mUNO targeting, we included St-PGA-DOX as an untargeted control. We employed ^1^H NMR and UV-Vis analyses to evaluate the chemical identity of nanoconjugates (Fig. S10A, B).

OximUNO displayed DOX and mUNO loadings of ∼10% and ∼4% in weight, respectively, corresponding to around four DOX and seven mUNO molecules for every OximUNO. OximUNO exhibited a size of ∼40 nm and a highly negative surface charge of -40 mV (Fig. 3B, C). We obtained similar DOX loading, size by DLS, and surface charge values for St-PGA-DOX (Fig. 3B, C).

The pH-sensitive hydrazone linker and the intrinsic biodegradability of St-PGA by lysosomal protease cathepsin B are expected to secure DOX release from OximUNO after cell internalisation^51^. Hence, we studied DOX release kinetics from OximUNO in the presence of acidic pH (pH 5) and cathepsin B using liquid chromatography-mass spectrometry (LC-MS, Fig. S11A-G). As we aimed for the i.p. administration of OximUNO, we assessed DOX release in intraperitoneal fluid (i.p. fluid) (Fig. 3D). At pH 5, we observed a sustained DOX release in the first 8 h (reaching a plateau at 15%), thereby demonstrating the suitability for endo-lysosomal drug delivery. DOX release in the presence of cathepsin B displayed comparable values in the first 8 h (∼13%), followed by a plateau and a reduced rate in the following hours (∼13% cumulative release at 72 h). Importantly, OximUNO exhibited negligible drug release in both physiological conditions evaluated (PBS and i.p. fluid) (Fig. 3D).

We next evaluated the in vitro cytotoxicity of OximUNO and St-PGA-DOX in primary human blood monocyte-derived M2, M0 and M1 macrophages. Since the in vivo concentration that provided optimal CD206^+^ TAM targeting with minimal hepatic accumulation was 30 µM in OG, here we focused our interest on conjugates at 33 µM of DOX. Our previous studies comparing other mUNO-targeted vs. untargeted polymeric nanosystems^44^ demonstrated that the highest targeted uptake in primary M2 macrophages occurred after an interval of 10 to 30 min. For this reason, we used an incubation time of 15 min for these experiments.

Importantly, in M2 macrophages, OximUNO displayed a significantly higher toxicity than DOX and St-PGA-DOX (Fig. 3E). Since CD206 is expressed on macrophage populations in other organs, although at lower levels than for M2 TAMs, we also evaluated the effect on non-polarised, “M0”, macrophages, which we previously showed express CD206 at levels lower than in M2 macrophages^42^. OximUNO revealed a moderate toxicity to M0 macrophages (Fig. 3F), but lower than the toxicity to M2. Despite this moderate toxicity, as St-PGA-OG-mUNO did not target the lung, liver, or spleen (Fig. S6), we expect OximUNO not to affect the macrophage populations of those organs. St-PGA-DOX showed its highest toxicity in M1 macrophages (Fig. 3G). We speculate that here, the phagocytic activity, known to be highest for M1 macrophages^52,53^, governs the uptake and also explains the 60% cell viability observed for OximUNO (Fig. 3G). In vivo, St-PGA-OG-mUNO did not target M1 TAMs (Fig. 2C, I), hence, we do not expect OximUNO to affect this population. Free DOX only displayed toxicity in M2 macrophages at 100 µM (Fig. 3E).

These results provide evidence that OximUNO displayed increased toxicity towards M2 macrophages when compared to St-PGA-DOX or DOX alone.

We also evaluated the hepatic and renal safety profile of a single administration of OximUNO (at doses corresponding to 2 mg/kg and 4 mg/kg of DOX) by analysing creatinine (Crea) and alanine aminotransferase (ALAT) levels 48 h after i.p. administration in healthy mice (Fig. 3H). These doses did not induce toxic levels of Crea or ALAT compared to the values reported in the literature^54^ or the reference values for the female Balb/c reported in the Mouse Phenome Database by The Jackson Laboratory^55^ or Charles River facilities^56^. However, increased ALAT levels with the higher dose, prompted the selection of the OximUNO dose corresponding to 2 mg/kg of DOX for further in vivo studies.

In summary, the conjugation of mUNO and DOX to the St-PGA backbone to yield OximUNO, enhanced the in vitro efficacy of DOX towards M2 macrophages with no in vivo renal or hepatic toxicity observed.

### OximUNO treatment of orthotopic TNBC depletes CD206^+^ TAMs, inhibits tumour progression and attenuates immunosuppression

The findings of the in vivo homing and in vitro cytotoxicity studies supported the subsequent evaluation of OximUNO in the orthotopic TNBC model. When tumours reached 25 mm^3^, we treated mice with i.p. injections of OximUNO, St-PGA-DOX, or DOX, at 2 mg/kg of DOX every other day for eighteen days. Encouragingly, OximUNO treatment significantly reduced primary tumour volume growth kinetics (Fig. 4A, red line) compared to DOX, St-PGA-DOX, and PBS. Furthermore, only the OximUNO treatment significantly reduced final tumour weight (Fig. 4B) compared to the untreated group. We assigned this encouraging therapeutic effect to mUNO-mediated targeting, as animals treated with the untargeted St-PGA-DOX possessed tumour volumes (Fig. 4A, blue line) similar to the PBS group (Fig. 4A, black line). Furthermore, OximUNO treatment did not affect mouse bodyweight, whereas treatment with DOX induced a significant decrease in mouse bodyweight starting from day twenty-one p.i. until the end of the treatment (Fig. 4C).

**Figure 4.**
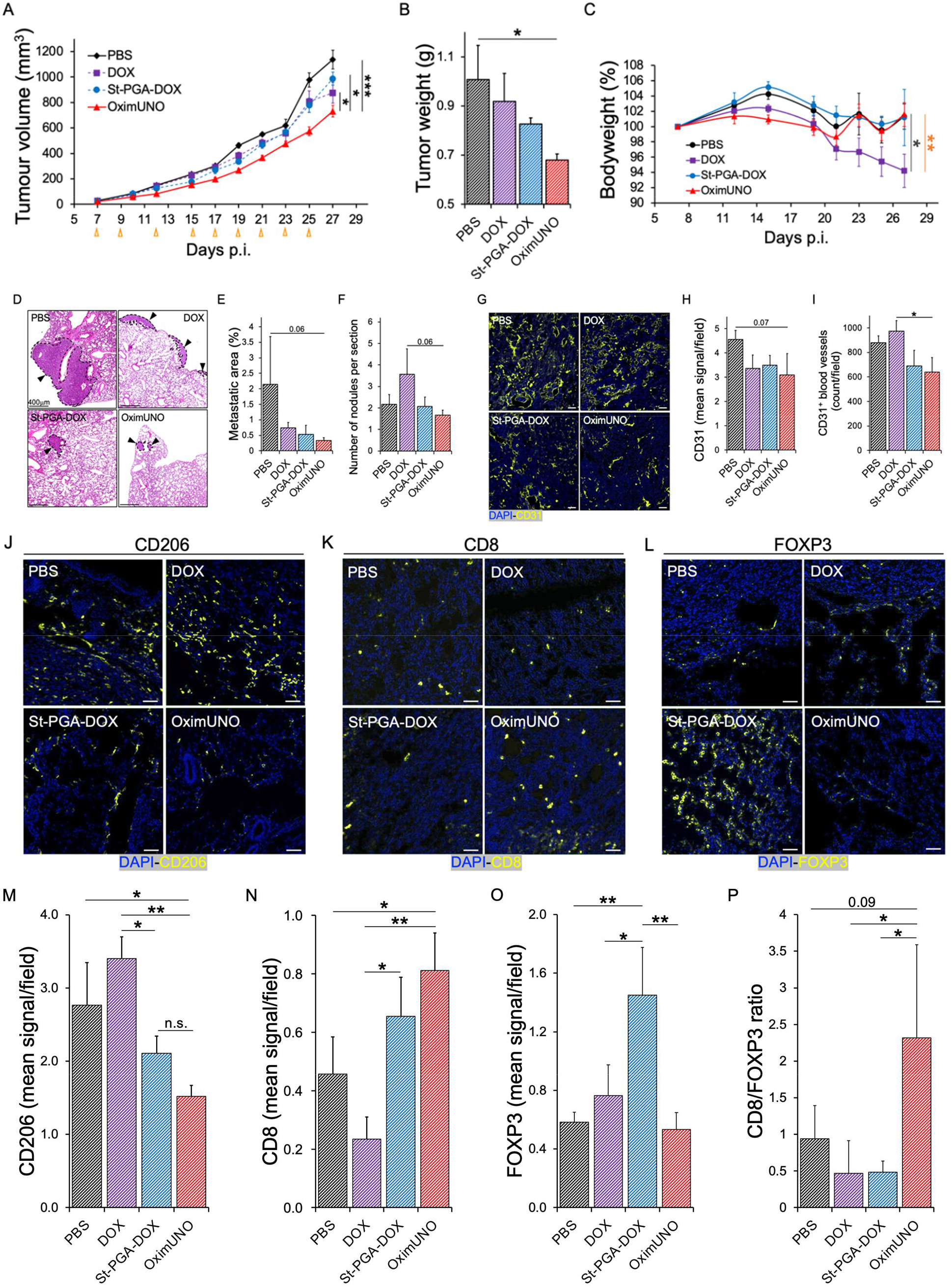
OximUNO treatment reduces primary tumour growth and pulmonary metastases and alleviates immunosuppression. Treatment with OximUNO, St-PGA-DOX, or DOX at 2 mg/kg of DOX in mice bearing orthotopic TNBC tumours (N=5). I.p. injections began when tumours reached 25 mm^3^ and were performed every other day to give a total of nine injections. (**A**) Primary tumour volume progression during treatment. Orange arrows indicate injection days. (**B**) Primary tumour weight at the experimental endpoint, demonstrating a significantly smaller weight for OximUNO-treated mice (red bar) than other groups. (**C**) Mouse bodyweight analysis suggests the safety of OximUNO treatment (red line); meanwhile, DOX-treatment induced a significant reduction in bodyweight by the experimental endpoint (purple line). Dark grey * DOX vs. PBS, orange * DOX vs. St-PGA-DOX. (**D**) Representative H&E images showing pulmonary metastases for all groups (scale bars = 400 μm); OximUNO treatment associated with (**E**) the smallest metastatic area and (**F**) the lowest number of average nodules per lung. **(G)** Representative images showing the expression of CD31 and blood vessels. **(H)** Graph depicting the expression of CD31. **(I)** Graph depicting the CD31^+^ blood vessel count. CD31 expression and blood vessel count were calculated using ImageJ and five images per mouse per group for expression analysis and at least three images per mouse per group for blood vessel count. (**J-L**) Representative confocal microscopy images demonstrating the expression of (**J)** CD206, (**K**) CD8, and (**L**) FOXP3. Scale bars = 50 μm. (**M-P**) Quantification of confocal microscopy images for the expression of (**M**) CD206, (**N**) CD8, and (**O**) FOXP3. (**P**) Graph of CD8/FOXP3 expression ratio showing a shift in the immune profile. Quantification was performed using the ImageJ programme from at least three images per mouse and five mice per group. Error bars represent the SE of the mean.

Histological analysis of lungs from treated mice (Fig. S12 shows an H&E stain from a healthy lung for comparison) revealed that OximUNO also influenced the extent of pulmonary metastases (Fig. 4D), as it elicited the highest reduction in the metastatic lung area and nodule number (p=0.06 vs. PBS and p=0.06 vs. DOX, respectively (Fig. 4E, F)). Meanwhile, immunofluorescence (IF) microscopy revealed no significant changes in CD31 expression in tumours (Fig. 4G, H), but significantly fewer CD31^+^ structures in the OximUNO-treated mice compared to DOX-treated mice (Fig. 4G, I), suggesting that the reduction in nodule number in the OximUNO group (of Fig. 4F) may be mediated by the lower vascularisation in the primary tumour. Importantly, histological analysis revealed no cardiotoxicity in any treatment groups (Fig. S13). IF analysis revealed that only OximUNO significantly reduced the CD206 expression (assigned to CD206^+^ TAMs), compared to PBS (Fig. 4J, M). Interestingly, treatment with DOX upregulated CD206 expression (Fig. 4J, M), which agrees with previous reports that demonstrated an increase in the number of CD206^+^ TAMs following chemotherapy^24^.

Notably, only OximUNO treatment significantly increased CD8 expression (a marker of cytotoxic T cells (CTLs)) compared to PBS and DOX treatment (Fig. 4K, N). Unexpectedly, St-PGA-DOX treatment increased the expression of FOXP3, a marker for regulatory T cells (Tregs) (Fig. 4L, O). Analysis of the CD8/FOXP3 expression ratio revealed that OximUNO treatment resulted in a five-fold increase compared to St-PGA-DOX or DOX treatment (Fig. 4P), suggesting that OximUNO stimulated a shift in the immune landscape towards immunostimulation. Of note, in all cases, we normalised the quantification of marker expression using immunofluorescent images to the tissue area to account for different amounts of tissue in different images.

By targeting CD206^+^ TAMs with DOX via OximUNO treatment, we increased the efficacy and reduced the toxicity of DOX in the orthotopic TNBC model. Our results also suggest that the depletion of CD206^+^ TAMs by OximUNO elicited an immunostimulatory shift.

### OximUNO treatment of experimental TNBC metastasis reduces CD206^+^ TAMs number, tumour burden and attenuates immunosuppression

We next evaluated the effect of OximUNO on experimental TNBC metastasis using GFP-labelled 4T1 cells. We treated mice every other day with i.p. injections of OximUNO, St-PGA-DOX, or DOX, starting from day four p.i. and sacrificed mice on day eighteen p.i. Analysis of whole lung fluorescence in the green channel revealed that OximUNO treatment induced the lowest GFP fluorescence, indicating a lower level of pulmonary metastases (Fig. 5A). Representative macroscopic images also provided evidence for a reduction in metastases (Fig. 5B). Confocal fluorescence microscopy of lungs confirmed the trend observed with whole lung fluorescence, showing fewer GFP fluorescent nodules in the OximUNO-treated group (Fig. 5C). Furthermore, histological analysis of lungs displayed the lowest number of pulmonary nodules for OximUNO-treated mice (Fig. 5D). Mice treated with the untargeted St-PGA-DOX and free DOX showed a significant decrease in bodyweight, resulting in a 19% (Fig. 5E, blue line) and 17% loss (Fig. 5E, purple line), respectively; meanwhile, OximUNO-treated mice displayed lower bodyweight loss (Fig. 5E, red line).

**Figure 5.**
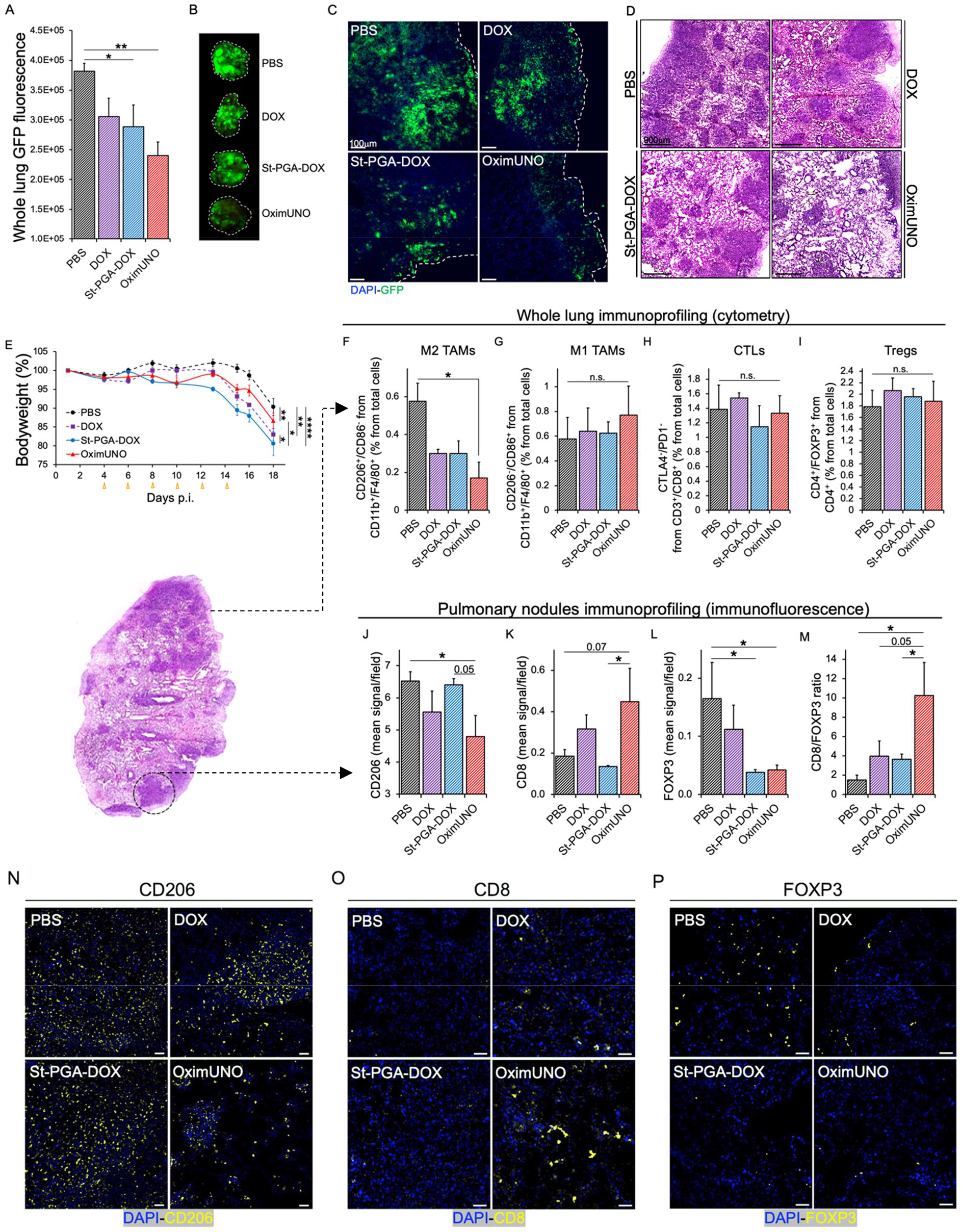
OximUNO treatment in experimental metastasis of TNBC significantly reduces CD206^+^ TAM number and tumour burden and alleviates immunosuppression. Treatment with OximUNO, St-PGA-DOX, or DOX at 2 mg/kg of DOX in the experimental metastasis of TNBC model, created using GFP-labelled 4T1 cells (N=6). I.p. injections began on day four p.i. and were performed every other day to give a total of six injections. (**A**) Quantification of whole lung GFP fluorescence at the experimental endpoint using the ImageJ programme (N=6). (**B**) Representative macroscopic photographs of GFP fluorescence in the lungs. (**C**) Representative confocal microscopy images of GFP expression, scale bars = 100 μm. (**D**) Representative H&E images showing pulmonary metastases for all groups (scale bars = 900 μm). (**E**) Mouse bodyweight analysis, demonstrating significantly lower bodyweight lost with OximUNO (red line) compared with St-PGA-DOX-treated mice (blue line) and DOX-treated mice (purple dotted line). Orange arrows indicate injection days. (**F-I**) FC analysis on three right lungs per group. (**F**) M2 TAMs (CD206^+^), (**G**) M1 TAMs, (**H**) CTLs and (**I**) Tregs. (**J-M**) IF analysis on the pulmonary tumour nodules to detect the expression of (**J**) CD206, (**K**) CD8, and (**L**) FOXP3. (**M**) Graph showing CD8/FOXP3 expression ratio. IF images quantified using the ImageJ programme from at least five images per mouse and three mice per group. (**N-P**) Representative confocal microscopy images for (**N**) CD206, (**O**) CD8, and (**P**) FOXP3. Scale bars = 50 μm. Error bars represent the SE of the mean.

We next employed flow cytometry (FC) to analyse the effect of different treatments on the immune cell populations in whole lungs. This analysis demonstrated that OximUNO treatment significantly lowered the percentage of M2 TAMs (CD206^+^) (Fig. 5F) but did not significantly impact the percentage of M1 TAMs, CTLs, or Tregs (Fig. 5G-I). We observed the same trend when we expressed these populations as total cell counts (Fig. S14-S17).

To evaluate if OximUNO affected CD206^+^ macrophages other than M2 TAMs, we analysed the state of splenic macrophages from this treatment study using FC. This analysis revealed no significant differences in the CD206/CD86 populations between the OximUNO-treated mice and PBS-treated mice (Fig. S18A-C).

While FC analysis informs on the immune status of the whole lung, it does not provide specific information regarding the TME. To characterise the immune landscape of the TME, we next analysed the expression of markers for TAMs, CTLs, and Tregs in pulmonary nodules using IF. This analysis revealed significantly lower CD206 expression in OximUNO-treated mice than PBS-treated mice (Fig. 5J, N), providing evidence for a robust reduction in the number of CD206^+^ TAMs in the TME. Importantly, and similarly to OximUNO treatment in the orthotopic TNBC mouse model, OximUNO elicited the highest expression of CD8 (Fig. 5K, O). OximUNO and St-PGA-DOX treated mice demonstrated significantly lower lung FOXP3 expression than PBS- and DOX-treated mice (Fig. 5L, P). OximUNO-treated mice displayed between a two- and three-times higher CD8/FOXP3 expression ratio than St-PGA-DOX and DOX, and nearly seven-times higher than PBS (Fig. 5M). Therefore, our IF analysis in the pulmonary tumour nodules suggested that OximUNO triggered a shift in the immune profile of the TME towards immunostimulation.

By targeting DOX to CD206^+^ TAMs in experimental TNBC metastases, we increased the efficacy and reduced the toxicity of DOX, as OximUNO treatment associated with the presence of fewer pulmonary tumour lesions and less bodyweight loss when compared to treatment with untargeted St-PGA-DOX and DOX. Our results suggest that the observed therapeutic effect derived from CD206^+^ TAM depletion, which elicited an immunological shift in the TME.

## DISCUSSION

To date, TNBC remains an aggressive breast cancer subtype^3^ with few treatment options, with conventional chemotherapy representing the current standard of care^20^. ICIs for TNBC have provided only modest improvements in complete response and progression-free survival in a small subset of TNBC patients^9,12,15,16^. Targeting TAMs can potentiate ICIs and other modalities and, therefore, represents an intense area of study^57–61;^ however, TAMs represent a diverse population^62–64^, and which TAM subtype to target remains under investigation.

Promising TAM-focused interventions under clinical evaluation include antibody-mediated depletion of TREM2-expressing TAMs (clinical trial identifier: NCT04691375). Antibody blockade of Clever-1 on M2 TAMs stimulated an M2→M1 switch in TNBC models (4T1) and synergised with the PD-1 blockade^65^. Appealing studies have used anti-CD163 antibodies to target TAMs^66^ by decorating DOX-carrying liposomes with anti-CD163, to deplete TAMs and potentiate ICIs in melanoma. Given our data comparing the tumour penetration of an anti-PDL1 antibody vs. St-PGA-OG-mUNO, anti-CD163 systems may also display lower tumour accumulation than St-PGA-OG-mUNO and OximUNO. Strategies targeting generic TAM markers such as CSF1R and CCR2 have shown side effects and limited efficacy.

Motivated by the preponderance of the mannose receptor in tumourigenic/metastatic TAMs in breast cancer^67–69^, here, we set out to deplete CD206^+^ TAMs in aggressive and metastatic TNBC models and study the consequences on the progression and immunosuppressive state of the tumour. To target CD206, a CD206-binding nanobody was developed by Ginderachter et al.^70^ which showed homing to CD206^+^ TAMs in in vivo models of lung and breast cancers^70^. Navidea Inc. engineered a mannosylated compound^71^ (Manocept™), that forms part of the FDA-approved contrast agent Lymphoseek®. Unfortunately, mannose-based ligands have other binding partners besides CD206, including CD209 in intestinal and genital tissues^45^, and can target dendritic cells^46^. Riptide Inc. also designed a peptide (RP-182) that binds to CD206; however, the peptide also binds to RelB, Sirp-α and CD47^72^.

We recently identified and described a short peptide called mUNO (sequence: CSPGAK) that targets mouse^41^ and human CD206^43^ at a different binding site than for mannose on CD206^43^. We identified mUNO from an in vivo screen using a peptide library in mice bearing metastatic breast cancer; we subsequently described how mUNO homed to CD206^+^ TAMs in other solid tumour models^41,73^ and in early-stage models of TNBC^42^ displaying low hepatic accumulation. We envisioned that conjugating mUNO to St-PGA would significantly enhance targeting through the avidity effect and increased plasma half-life^74^.

Compared to synthetic polymers such as N-(2-hydroxypropyl) methacrylamide, polypeptide-based nanocarriers show several benefits, including biodegradability, lower immunogenicity, and a lack of long term-accumulation, and the number of polypeptide-based constructs reaching clinical evaluation has significantly increased in recent years^75–77^. We employed St-PGA-based nanoconjugates with three linear chains (∼50 glutamic acids each) linked to a central core. Overall, the safety, lack of toxicity, and biodegradability of St-PGA meet FDA approval criteria^78^. A previous screen of PGA structures suggested that larger architectures enhanced plasma half-life and increased bioavailability through a higher hydrodynamic volume that reduces rapid renal clearance^47,79^. Of note, an extended plasma half-life will be advantageous when targeting the continuously replenished TAM cell type^80,81^.

St-PGA-OG-mUNO, a fluorescent counterpart of OximUNO, can be easily monitored by immunostaining for OG or detecting native OG fluorescence (as for the half-life study). Given weak DOX fluorescence and the inability to detect DOX with an antibody, we first designed St-PGA-OG-mUNO for validation purposes. We then exchanged OG for DOX to generate St-PGA-DOX-mUNO, referred to as “OximUNO”. Our studies demonstrated that St-PGA-OG-mUNO displayed a far greater plasma half-life and specificity to CD206^+^ TAMs than free mUNO and avoided CD86^+^ M1 TAMs and CD11c^+^ DCs, an important fact since M1 TAMs display anti-tumourigenic activity^25^, and CD11c^+^ DCs participate in antigen presentation^82^. In line with these observations, the computational analysis indicated that mUNO peptides are available to a receptor and sweep a vast space (130°) around PGA. Altogether this data demonstrates the benefit of conjugating mUNO to St-PGA. While previous studies have reported the St-PGA nanocarrier^47,78^ and the mUNO targeting peptide^42^, this work represents the first design of a peptide-targeted St-PGA nanosystem. Regarding the administration route of peptide-guided St-PGA nanosystems, in the future we also wish to evaluate the i.v. route, which, barring the case of intraperitoneal chemotherapy, represents a more clinically translatable route to deliver cancer therapies.

In the OximUNO system, drug release studies revealed only 15% DOX release, which agrees with our previous studies^48,50^ but suggests room for improvement, which may come from using longer polymer-drug linkers such as EMCH (N-ε-maleimidocaproic acid hydrazide) moiety^48,50^ or from the use of external triggers^83–85^. Unexpectedly, we failed to observe a significant increase in DOX release in the presence of cathepsin B with respect to the hydrolytic conditions; we hypothesise that the nanoconjugate conformation slows down proteolytic degradation, hampering in vitro quantification within the studied timeframe^47^.

Our in vivo efficacy studies showed that, strikingly, the sole depletion of CD206^+^ TAMs with OximUNO alleviated tumoural immunosuppression and reduced dissemination and growth, confirming the pro-tumoural and immunosuppressive roles assigned to CD206^+^ TAMs in the literature and reaffirming the importance of targeting this particular TAM subset. Additionally, the observed reduction in the number of CD206^+^ TAMs and CD31^+^ structures for OximUNO agrees with the established angiogenic role of CD206^+^ TAMs^24^.

From a safety point of view, we found that the OximUNO nano-formulation of DOX had the least negative impact on mouse bodyweight compared to free DOX or the untargeted nano-formulation St-PGA-DOX. Additionally, OximUNO did not alter Crea or ALAT levels, indicating the absence of acute hepatic or renal toxicity. Our data suggest that the signal observed in the kidneys for St-PGA-OG-mUNO (consistent with the previously reported excretion of St-PGA^47,78^) did not translate into acute renal toxicity for OximUNO. These are relevant findings as DOX induces cell death and tissue damage not only in the heart but also in the liver and kidneys^86^. OximUNO displayed moderate toxicity to un-polarised macrophages in vitro; however, OximUNO did not affect or alter the macrophage populations of the spleen in vivo, in agreement with the absence of spleen targeting we observed for St-PGA-OG-mUNO.

Most preclinical studies evaluating the effect of M2 TAM targeted monotherapy in the 4T1 mouse model have either not shown efficacy on secondary tumours^87,88^, a lack of efficacy in primary tumours or metastases in the case of anti-CLEVER-1^65^, or a pro-metastatic effect in the case of anti-CSF1R^89^. Hence, along with anti-MARCO therapy^90^, OximUNO constitutes one of the few reports of an M2 TAM targeted monotherapy affecting both primary and secondary tumours in the 4T1 mouse model.

Beyond TAM depletion, we show that St-PGA-mUNO represents an attractive platform to carry additional therapeutic payloads other than DOX, which could include M2→M1 polarising agents such as TLR7 agonists^44,91^, beta-emitting radiotherapeutic agents such as dodecanetetraacetic acid-chelated ^177^Lu ^92^, or photosensitisers used in photodynamic therapy^83–85^. We also envisage the combination of TAM-depletion via OximUNO administration together with current chemotherapy regimens to prevent dissemination and relapse, or the use of OximUNO prior to surgery, i.e., as neoadjuvant chemotherapy

Taking OximUNO as a proof-of-concept, our data support the peptide-targeted St-PGA design reported here as a new targeted nanosystem that could target other receptors by exchanging the targeting peptide.

## MATERIALS AND METHODS

### Reagents and solutions

The peptides mUNO (sequence: CSPGAK-COOH) and FAM-mUNO (FAM-Ahx-CSPGAK-COOH) were purchased from TAG Copenhagen and doxorubicin (DOX) from Sigma-Aldrich. St-PGA was kindly provided by Polypeptide Therapeutic Solution S.L. (PTS, Valencia, Spain). See the Supplementary Information for information on all other reagents and solutions.

<u>Mayer’s haematoxylin solution</u> was prepared by dissolving 5 g of aluminium potassium sulphate dodecahydrate (Merck Millipore, cat. 1010421000) in 100 mL of water, and adding 1 g of haematoxylin (Merck, cat. H9627). After complete dissolution, 0.02 g of sodium iodide (Merck, cat. 1065230100) was added and completely dissolved. Then, 2 mL of acetic acid (Sigma-Aldrich, cat. 33209) was added, and then the solution was boiled and then cooled. Once ready to use, the solution was filtered using a 0.45 μm filter.

<u>Eosin (5%) solution</u> was prepared by dissolving 0.5 g of Eosin Y (Sigma-Aldrich, cat. 230251) in 99 mL water/1 mL acetic acid.

### Cell culture and experimental animals

4T1 cells were purchased from ATCC, and 4T1-GFP cells were a gift from Ruoslahti laboratory (Sanford-Burnham-Prebys Medical Discovery Institute, La Jolla, USA). 4T1 and 4T1-GFP cells were cultured in RPMI-1640 medium (Gibco by Life Technologies, cat. 72400-021) supplemented with 10% v/v foetal bovine serum (FBS, Capricorn Scientific, cat. FBS-11A) and 100 IU/mL penicillin/streptomycin (Capricorn Scientific, cat. PS-B) at 37 °C in the presence of 5% CO_2_. For all animal experiments, 8-12-week-old female Balb/c mice were used. Animal experiment protocols were approved by the Estonian Ministry of Agriculture (Project #159). All methods were performed in accordance with existing guidelines and regulations.

### Tumour models

Two tumour models were used for homing studies: the orthotopic TNBC model, where 1×10^6^ 4T1 cells in 50 µL of phosphate-buffered saline (PBS, Lonza, cat. 17-512F) were subcutaneously (s.c.) injected into the fourth mammary fat pad, and the experimental metastasis of TNBC model, where 5×10^5^ 4T1 cells in 100 µL of PBS were injected i.v. into Balb/c mice. Two tumour models were used for treatment studies: the orthotopic TNBC model where 5×10^4^ 4T1 cells in 50 µL of PBS were injected s.c. into fourth mammary fat pad; and the experimental metastasis of TNBC model where 2×10^5^ 4T1-GFP cells in 100 µL of PBS were i.v. injected.

### Nanoconjugate synthesis and characterisation

In vivo homing studies used St-PGA-OG and St-PGA-OG-mUNO, while in vitro cytotoxicity and in vivo treatment studies used St-PGA-DOX and St-PGA-DOX-mUNO (“OximUNO”). Detailed synthetic procedures for single nanoconjugates can be found in Supplementary Information.

### Physico-chemical characterisation methods

<u>Nuclear magnetic resonance (NMR) spectroscopy:</u> NMR spectra were recorded at 27 °C (300 K) on a 300 UltrashieldTM from Bruker. Data were processed with Mestrenova software. Sample solutions were prepared at the desired concentration in D_2_O or D_2_O supplemented with NaHCO_3_ (0.5 M).

<u>UV-Visible (UV-Vis) analysis:</u> UV-Vis measurements were performed using JASCO V-630 spectrophotometer at 25 °C with 1 cm quartz cells and a spectral bandwidth of 0.5 nm. Spectra analysis was recorded three times in the range of 200–700 nm.

<u>Fluorescence analysis:</u> Fluorescence analysis was performed using a JASCO FP-6500 spectrofluorimeter at 25 °C with 1 cm quartz cells

<u>Dynamic Light Scattering (DLS):</u> Size measurements were performed using a Malvern ZetasizerNano ZS instrument, supported by a 532 nm laser at a fixed scattering angle of 173°. Nanoconjugate solutions (0.1 mg/mL) were freshly prepared in PBS (10 mM phosphate, 150 mM NaCl), filtered through a 0.45 μm cellulose membrane filter, and measured. Size distribution was measured (diameter, nm) for each polymer in triplicate. Automatic optimisation of beam focusing and attenuation was applied for each sample.

<u>Zeta potential measurements:</u> Zeta potential measurements were performed at 25 °C using a Malvern ZetasizerNano ZS instrument, equipped with a 532 nm laser using disposable folded capillary cells, provided by Malvern Instruments Ltd. Nanoconjugate solutions (0.1 mg/mL) were freshly prepared in 1 mM KCl. Solutions were filtered through a 0.45 μm cellulose membrane filter. Zeta potential was measured for each sample per triplicate.

### Molecular dynamics simulations

Molecular dynamics (MD) simulations of PGA chains, and mUNO peptide were carried out using the ff19SB force field^93^ in the Amber20 MD engine^94^. The nanoconjugate system was neutralised using Na+ ions and hydrated to account for a total of ∼920,000 atoms (∼300,000 TIP3P water molecules) in a truncated octahedral box. A hydrogen mass repartitioning strategy was applied on the resulting topology, allowing us a 4 fs integration time step^95^. Standard minimisation and equilibration protocols were used to reach 300 K and 1 atm., followed by 50 ns of production MD run. The simulations were run under the NVT ensemble (constant number of particles, volume, and temperature through Berendsen thermostat^96^), considering periodic boundary conditions. The SHAKE algorithm was used to fix hydrogen atoms^97^. The non-bound cut-off value was set to Angstrom. The central moiety was parametrised using the recommended protocol for the Amber force field. It was necessary to introduce amide bond, angle and dihedral terms using the ParmEd module to establish the bond of the central molecule to the PGA chains.

### Tumour homing studies

Tumours were induced as described in the tumour model section. Tumour homing studies were performed on mice bearing orthotopic TNBC or experimental metastasis of TNBC. Ten days p.i. of the orthotopic TNBC or the experimental metastasis of TNBC model, mice were i.p. injected with St-PGA-OG-mUNO (0.41 mg/0.5mL of PBS) or St-PGA-OG (0.35 mg/0.5mL of PBS) (corresponding to 15 nmoles of OG, fluorescence measured by UV-Vis). The homing of a higher dose of St-PGA-PGA-mUNO (0.82 mg/0.5mL of PBS) or St-PGA-OG (0.7 mg/0.5mL of PBS) (corresponding to 30 nmoles of OG) was also analysed and compared with the homing of FAM-mUNO (30 nmoles/0.5mL of PBS). In every case, nanoconjugates or free peptide were circulated for 6 h, after which time, mice were sacrificed by anaesthetic overdose followed by cervical dislocation. Organs and tumours were collected and fixed in cold 4% w/v paraformaldehyde (PFA) in PBS at +4 °C for 24 h, washed in PBS at room temperature for 1 h and cryoprotected in 15% w/v sucrose (Sigma Life Science, cat. S9378) followed by 30% w/v sucrose at 4 °C overnight. Cryoprotected and fixed tissues were frozen in OCT (Optimal Cutting Temperature, Leica, cat. 14020108926), cryosectioned at 10 µm on Superfrost+ slides (Thermo Fisher, cat. J1800AMNZ) and stored at -20 °C. Immunofluorescent staining was performed as described earlier^42^. OG was detected using rabbit anti-FITC/Oregon Green (dilution 1/100, Invitrogen by Thermo Fisher Scientific, cat. A889) and Alexa Fluor® 647 goat anti-rabbit antibody (dilution 1/250, Invitrogen by Thermo Fisher Scientific, cat. A21245). CD206 was detected using rat anti-mouse CD206 (dilution 1/150, Bio-Rad, cat. MCA2235GA) and Alexa Fluor® 546 goat anti-rat antibody (dilution 1/250, life technologies, cat. A11081). CD86 was detected using rat anti-mouse CD86 (dilution 1/100, BioLegend, cat. 105001) and Alexa Fluor® 546 goat anti-rat secondary antibody (dilution 1/250). CD11c was detected using hamster anti-mouse CD11c antibody (dilution 1/75, BioLegend, cat. 117301) and Alexa Fluor® 546 goat anti-hamster secondary antibody (dilution 1/200, life technologies, cat. A21111) Slides were counterstained using 4′,6-diamidino-2-phenylindole (DAPI, 1 μg/mL in PBS, Sigma-Aldrich, cat. D9542-5MG). Coverslips were mounted using mounting medium (Fluoromount-G™ Electron Microscopy Sciences, cat. 17984-25), and sections were imaged using Zeiss confocal microscope (Zeiss LSM-710) and 20x objective. The colocalisation analysis shown in Fig. 2M-P, between the FAM or OG channel and the CD206 channel was carried out using the “Coloc2” plugin in the Fiji programme and selecting the “Pearson’s R value (no threshold)” coefficient. The colocalisation values were obtained from at least three representative images per mouse per group and their average and standard error were plotted. The OG/FAM mean signal per CD206^+^ cell analysis was measured using the ImageJ programme, taking the mean OG/FAM signal, and dividing it with the number of CD206^+^ cells. Average values were obtained from four images per mouse. N=3 for orthotopic TNBC and N=2 for the homing in experimental metastasis of TNBC.

### Analysis of tumour and liver leakiness

Endogenous IgG immunostaining of orthotopic 4T1 tumours and livers was performed following the same IF protocol as described above to assess leakiness. Endogenous IgG was detected using Alexa Fluor® 647 goat anti-mouse antibody (dilution 1/200, Invitrogen by Thermo Fisher Scientific, cat. A21235) and slides were counterstained with DAPI (1 μg/mL in PBS). The coverslips were mounted, and sections were imaged using Zeiss confocal microscope and 20x objective (N=3 tumours).

### PDL1 expression analysis in orthotopic TNBC tumours

The assessment of PDL1 expression in orthotopic 4T1 tumours followed the IF protocol described above. PDL1 was detected using rat anti-mouse PDL1 (dilution 1/100, BioLegend, cat. 124302) as primary antibody and Alexa Fluor® 647 goat anti-rat (dilution 1/200, Invitrogen, cat. A21247) as the secondary antibody. Slides were counterstained with DAPI (1 μg/mL in PBS), mounted, and imaged using a Zeiss confocal microscope.

### Tumour homing of anti-PDL1 in orthotopic TNBC tumours

For the homing analysis with anti-PDL1, we injected 1×10^6^ 4T1 cells in 50 µL of PBS s.c. and ten days p.i., PDL1 antibody (5 mg/kg, rat anti-mouse, BioXcell, cat. BE0101) was injected i.v., circulated for 24 h after which time, mice were sacrificed, organs collected and fixed with PFA. Ten µm tissue sections were stained with Alexa Fluor® 647 goat anti-rat antibody (dilution 1/200), counterstained with DAPI (1 μg/mL in PBS), mounted, and imaged with a Zeiss confocal microscope.

### Plasma half-life evaluation for St-PGA-OG-mUNO

Plasma half-life studies were performed as previously described^42^. Briefly, healthy female Balb/c mice (N=3) were i.p. injected with St-PGA-OG-mUNO (0.41 mg/0.5mL of PBS, corresponding to 15 nmoles OG). Ten µL of blood was sampled at different timepoints (0, 5, 10, 15, 30, 60, 180, 360, and 1440 min) and mixed with 50 µL of PBS-Heparin solution. Blood samples were centrifuged to obtain plasma (300g for 5 min at room temperature) and OG fluorescence was read with a plate reader (FlexStation II Molecular Devices) at 480nm excitation/520nm emission.

### DOX release studies

LC-MS was implemented to determine free drug levels, stability, and drug release with OximUNO. The LC-MS system consisted of an ExionLC LC system and AB Sciex QTRAP 4500, a triple quadrupole ion trap hybrid equipped with a Turbo VTM electrospray ionisation source. DOX was detected with an internal standard method: 1 µg/mL of daunorubicin (DAU) was used as internal standard, where three calibration curves (in a range from 0.5 to 50 µg/mL DOX) were prepared and used for accurate analysis of DOX in the samples. Both DOX and DAU were detected with positive electrospray ionisation mode by following two mass transitions (544.2 m/z → 397 m/z and 544.2 m/z → 379 m/z for DOX, and 528 m/z → 363.1 m/z and 528 m/z → 321.3 m/z for DAU). The obtained LC-MS optimal conditions were as follows: flow rate 0.5 mL/min; mobile phase – 0.05 % trifluoroacetic acid with 70 % of acetonitrile; LiChrospher 100 C18 column (125×4.0 mm) (Merck); column temperature 40 °C, 10 µL injection volume.

### Stability study of OximUNO conjugate in PBS, pH 7.4

OximUNO was incubated in 10 mM dPBS at 37 °C at the concentration of 3 mg/mL and with 3 µg/mL of DAU. 100 µL aliquots were collected at defined time points (0, 1, 2, 5, 24, 48, 72 h), extracted with 3×250 µL chloroform, and mixed by vortexing for 5 min. Organic phases from all three chloroform extracts were collected in one tube, evaporated using speed vacuum, and stored at -20 °C. On the day of analysis, dried samples were reconstituted in 300 µL of methanol (LC-MS grade), vortexed for 5 min and centrifuged for 5 min at 30,437g. Supernatants were filtered through a 0.45 µm filter and subjected to LC-MS analysis.

### Stability study of OximUNO in the i.p. fluid

I.p. fluid was collected from healthy 8-12-week old Balb/c female mice as performed in REF^98^ by collecting the supernatant and discarding the pellet after the centrifugation step. A working solution containing 3 mg/mL of OximUNO and 3 µg/mL of DAU in i.p. fluid was incubated at 37 °C. 50 µL aliquots were collected at scheduled time points (0, 2, 5, 7, and 24 h). Samples were then diluted with 100 µL of methanol, sonicated to dissolve DOX, and injected into the LC-MS after filtration through a 0.45 µm filter.

### Cathepsin B release kinetic studies

Cathepsin B (5 IU) was activated in 2 mM EDTA, 5 mM DTT, and 20 mM CH_3_COONa buffer and incubated at 37 °C for 15 min. In a separate tube, a solution containing 3 mg/mL OximUNO and 3 µg/mL of DAU was prepared with 20 mM CH_3_COONa and incubated at 37 °C for 15 min. The two solutions were then combined to produce a reaction solution that was incubated at 37 °C. 100 µL aliquots were collected at scheduled time points (0, 1, 2, 5, 8, 24, 48, 72 h), and after the addition of 900 µL of dPBS (to adjust the pH level to 7.4), free DOX and DAU were extracted with 2.5 mL of CHCl_3_ three times. Samples were processed as described under “Stability Study of OximUNO conjugate in PBS, pH 7.4”. After CHCl_3_ evaporation, samples were reconstituted with 300 µL of methanol, filtered through a 0.45 µm filter and subjected to LC-MS analysis. A blank solution was prepared with the same components as the sample solution but without cathepsin B and used as a control sample.

### In vitro cytotoxicity assay

Human peripheral blood mononuclear cells (PBMC) were purified from human blood buffy coat using Ficoll Paque Plus (GE Healthcare, cat. 17-1440-02) reagent and CD14^+^ microbeads (MACS Miltenyi Biotec, cat. 130-050-201) as previously described^42^. 1.2×10^5^ cells in 50 μL of RPMI-1640 medium were seeded on an FBS-coated 96-well plate. To obtain M0 macrophages, 50 μL of macrophage colony stimulating factor (M-CSF) (100 ng/mL, BioLegend, cat. 574802) was added and replenished every other day for four days by substituting the half of the medium with fresh medium containing M-CSF. To obtain optimal macrophage attachment and M2 polarisation, 50 μL of interleukin-4 (IL-4, 50 ng/mL, BioLegend, cat. 574002) and M-CSF (100 ng/mL) mixture was added to the wells. The medium was replenished by substituting half of the medium with fresh medium containing IL-4 and M-CSF every other day for six days. To obtain M1 macrophages, monocytes were incubated with M-CSF (100 ng/mL) for six days, replenishing every other day with fresh medium containing M-CSF and on day six, 50 μL of M-CSF, lipopolysaccharide (LPS, 100 ng/mL, Sigma Aldrich, cat. L4391) and interferon-γ (IFN-γ, 20 ng/mL, BioLegend, cat. 570202) was added and incubated overnight. On day seven for M2 and M1 macrophages or day four for M0 macrophages, cells were incubated for 15 min at 37 °C with OximUNO, St-PGA-DOX, DOX in medium, or free medium as a control (N=3 wells per group). Concentrations used were calculated based on DOX: 33μM and 100μM. (Of note, the dose of OximUNO used for the 33 µM DOX experiments shown in Figure 3E corresponds to the same dose of OximUNO used for both in vivo treatment studies. In vivo, all treated groups received injections containing 2 mg/kg of DOX, which, assuming the dilution in mouse blood, corresponds to a DOX concentration of 33 µM). After incubation, wells were washed, fresh medium added, and cells incubated for 48 h at 37 °C. After 48 h, 10 μL of 3-(4,5-dimethylthiazol-2-yl)-2,5-diphenyltetrazolium bromide (MTT, concentration 5 mg/mL, Invitrogen, cat. M6494) in PBS was added to each well containing culture medium and incubated for 2.5 h at 37 °C. Medium containing MTT was then removed without removing formed crystals, and 100 μL of isopropanol was added to each well to dissolve crystals. Absorbance was read at 580 nm using a plate reader (Tecan Sunrise) and the corresponding Magellan™ 7 programme.

### In vivo liver and kidney toxicology studies with OximUNO

Three healthy 12-week-old female Balb/c mice were i.p. injected once with OximUNO (0.704 mg/0.5mL PBS or 1.408mg/0.5mL) and circulated for 48 h. Then, mice were anesthetised, and blood collected through retro-orbital bleeding into Lithium Heparin tubes (BD Vacutainer, cat. 368494). Blood samples were centrifuged at 1800g for 15 min at +4 °C and 400 μL of plasma was collected for analysis. Samples were analysed in Tartu University Hospital using a Cobas 6000 IT-MW (Roche Diagnostics Gmbh) machine and reagents for creatinine (CREP2, cat. 03263991) and alanine aminotransferase (ALTLP, cat. 04467388).

### OximUNO treatment of orthotopic TNBC

5×10^4^ 4T1 cells in 50 μL of PBS were s.c. injected into the fourth mammary fat pad of 8-12-week-old female Balb/c mice. On day seven, mice were sorted into four groups by tumour volume measured using a digital calliper (Mitutoyo). Tumour volume was calculated based on the formula (W^2^ x L)/2, where W is the tumour’s width and L is the tumour’s length. The starting volume for each group was ∼25 mm^3^, and the number of mice in each group was five. The first i.p. injection of compounds was carried out on day seven, followed by an i.p. injection every other day; nine injections were performed in total. The dose of nanoconjugates was calculated based on DOX, 2 mg/kg per injection (DOX: 39.5 μg/0.5mL PBS; St-PGA-DOX: 476 μg/0.5mL PBS; OximUNO: 341 μg/0.5mL PBS) giving a cumulative dose of DOX of 18 mg/kg. Mouse bodyweight and tumour volumes were monitored every other day. The final injection was on day 25 and all mice were sacrificed on day 28. Tumour tissues were processed as described under “In vivo biodistribution studies”, and the lungs and hearts were embedded in paraffin and processed for haematoxylin and eosin (H&E) staining (described below). Tumours were immunostained as described above. CD206 was detected using rat anti-mouse CD206 (dilution 1/200), CD8 using rat anti-mouse CD8 (dilution 1/75 Biolegend, cat. number 100701), FOXP3 using rat anti-mouse FOXP3 (dilution 1/75, Biolegend, cat number 126401) as primary antibodies, Alexa Fluor® goat anti-rat 647 (dilution 1/300 for CD206 and 1/200 for CD8, FOXP3,) was used as a secondary antibody for all markers. Slides were counterstained with DAPI (1 μg/mL in PBS) and imaged using a Zeiss confocal microscope with a 10x objective. All five tumours from each group were included in the IF analysis and at least three images per mouse per group were included. Fluorescent signal intensity was calculated using the ImageJ programme; to account for different amounts of tissue in the different images, only the area containing tissue was selected and the “mean signal intensity” given by the programme taken (total integrated intensity divided by the selected area). For this analysis, at least three images per tumour were included.

### H&E staining in paraffin-embedded formalin-fixed tissues

For H&E staining, 2 μm sections were cut from paraffin-embedded blocks. Slides were warmed at 60 °C for 2 min before deparaffinising using xylene (3×2 min, 1×1 min) followed by 100% ethanol washes (3×1 min), 80% ethanol wash (1×1 min) followed by 1 min wash in water. Slides were first incubated with ST-1 HemaLast for 30 s, followed by ST-2 Haematoxylin for 5 min after which time, slides were washed in water for 2 min. Then, ST-3 Differentiator was added for 45 s, and slides were washed in water for 1 min. Next, ST-4 Bluing Agent was added (1 min), washed for 1 min in water followed by 1 min incubation in 80% ethanol, after which time, ST-5 Eosin was added and incubated for 1 min. For rehydration, incubations in 100% ethanol (2×30 s, 1×2 min) were carried out and finished with incubations in xylene (2×2 min). All washes were carried out in tap water. H&E stainings were performed in Tartu University Hospital by pathologists using Leica staining automat and ST Infinity H&E Staining System (Leica, cat. 38016998). Stained lung sections were scanned using a slide scanner (Leica SCN400) and 20x zoom. Images were analysed using the QuPath programme (version 0.1.2)^99^. Five levels ∼1 mm apart were used for each mouse to obtain comprehensive pulmonary metastases profile. Stained heart sections were also scanned using a slide scanner and analysed with the QuPath programme. Tartu University Hospital pathologists assessed cardiotoxicity in hearts and pulmonary metastases.

### Analysis of CD31 expression and blood vessel count

CD31 expression after treating orthotopic TNBC tumours with OximUNO, St-PGA-DOX, or DOX was detected using rat anti-mouse CD31 (dilution 1/100, BD Biosciences, cat. 553370) and Alexa Fluor® 546 goat anti-rat (dilution 1/200, Invitrogen, cat. A11081) was used as the secondary antibody. Slides were counterstained with DAPI (1 μg/mL in PBS) and imaged using Zeiss confocal microscope with a 10x objective. CD31 expression was calculated using ImageJ and mean signal per field as described under “OximUNO therapy in orthotopic TNBC”, including at least five images per mouse per group, N=5 mice per group. The blood vessel count was calculated from the same images using ImageJ as follows: the image was changed to an 8-bit image, threshold (Triangle algorithm with modifications to account for as much actual CD31 signal as possible) was added, and particles analysed. At least three images per mouse per group were included in the analysis, N=5 mice per group. Field size was 1.42 mm x 1.42 mm for all images.

### OximUNO treatment of experimental metastasis of TNBC

2×10^5^ 4T1 cells in 100 μL of PBS were i.v. injected into the tail vein of 8-12-week-old female Balb/c mice. Treatment with OximUNO, St-PGA-DOX, or DOX began on day four p.i.; each group comprised six mice. Doses of different compounds were calculated based on DOX (2 mg/kg): DOX: 39.5 μg/0.5mL PBS; St-PGA-DOX: 774.5 μg/0.5mL PBS; OximUNO: 704 μg/0.5mL PBS. Mouse bodyweight was monitored every other day. A total of six injections were carried out every other day. The final injection was on day 12, and all animals were sacrificed on day 18 using anaesthetic overdose and perfusion with PBS. Three right lungs from each group were analysed with flow cytometry (FC), and three full lungs and three left lungs from each group were frozen into blocks using OCT. Frozen lung tissues were cryosectioned as described earlier, fixed with cold 4% PFA (CD206) or acetone (for CD8 and FOXP3), and stained as described in the following section. Immunofluorescent stainings were performed using the same markers and antibodies as shown in the “OximUNO treatment in orthotopic TNBC” section.

### GFP staining and imaging

Six lungs from each group were frozen in OCT. Ten μm sections were cut, and slides were kept at -20 °C until ready to use. Slides were taken out of the freezer at least 30 min before staining. For staining, slides were fixed with 4% PFA for 10 min at room temperature, washed with PBS for 10 min at room temperature, counterstained using DAPI (1 μg/mL in PBS) for 5 min at room temperature, washed 3×4 min with PBS and finally mounted using mounting medium. Permeabilisation was not used in this step to improve GFP visualisation. GFP was visualised using its native fluorescence. Slides were imaged using Olympus confocal microscope (FV1200MPE) with a 10x objective.

### Macroscopic analysis of GFP Signal

Lungs from each group were imaged using Illumatool Bright Light System LT-9900 (LightTool’s Research) in the green channel to visualise the fluorescent signal macroscopically, and a photograph of each lung was taken. The total GFP signal of each lung was quantified using the ImageJ programme using the “IntDen” value.

### Flow cytometry analysis

Three mice were sacrificed using anaesthetic overdose, perfused with PBS and right lung tissues were placed in cold RPMI-1640 medium supplemented with 2% v/v FBS. Lungs were cut into small pieces on ice in a solution containing collagenase IV (160 U/mL, Gibco cat.17104019)/dispase (0.6 U/mL, Gibco, cat. 17105-041)/DNase I (15 U/mL; AppliChem, cat. A3778) mixture. To obtain a single-cell suspension, lung pieces were incubated in 10 mL of the same mixture at 37 °C on a rotating platform for 45-60 min, pipetting every 10 min to improve digestion. The cells were washed with 5 mL of RB (“running buffer”: 4 mL 0.5M EDTA, 100 mL v/v FBS in 1L of PBS), centrifuged (350g, 7 min, 4 °C), and red blood cells were lysed with 3 mL of ammonium-chloride-potassium lysing buffer (ACK) at room temperature. Ten mL of RB was added, cells were centrifuged and filtered using a 100 μm cell strainer (Falcon, cat. 352360). Cells were counted using the brightfield mode of LUNA™ Automated Cell counter (Logos Biosystems). Cells were collected in RB at a concentration of 5×10^6^/100μL, placed on a 96-well plate with conical bottom and incubated for 30 min in FcR-blocking 2.4G2 hybridoma medium at 4 °C. The cells were then stained for either macrophage or T cell markers for 25-45 min in the dark at +4 °C, centrifuged and washed twice with RB. The antibodies used are listed in Table 2. For intracellular staining of T cells, cells were fixed using eBioscience™ FOXP3/Transcription Factor Staining Buffer Set (Thermo Fisher, cat. 00-5523-00) according to the protocol provided. Cells were stained for 25-45 min in the dark at room temperature following permeabilisation and washed twice using RB. All cells were collected in 150 μL of RB, filtered through a 70 μm filter (Share Group Limited) and 150 μL of RB was used to wash the filter. BD LSRFortessa Flow Cytometer and FCS Express 7 Flow (De Novo Software) were used for analysis.

**Table 2.** Antibodies used in FC analysis: macrophage and T cell markers.

|  | Antibody | Dilution, company |
| --- | --- | --- |
| Macrophage markers | PerCP/Cyanine5.5 anti-mouse CD206 (MMR) | 1/200, BioLegend, clone C068C2, cat. 141715 |
|  | PE anti-mouse CD86 | 1/400, BioLegend, clone PO3, cat. 105105 |
|  | PE/Cyanine7 anti-mouse F4/80 | 1/200, BioLegend, clone BM8, cat. 123114 |
|  | PE/Dazzle™ 594 anti-mouse/human CD11b | 1/ 800, BioLegend, clone M1/70, cat. 101255 |
|  | eBioscience™ Fixable Viability Due eFluor™ 506 | 1/800, Thermo Fisher Scientific, cat. 65-0866-18 |
| T cell markers | Brilliant Violet 570™ anti-mouse CD4 | 1/400, BioLegend, clone RM4-5, cat. 100542 |
|  | Brilliant Violet 605™ anti-mouse CD8a | 1/400, BioLegend, clone 53-6.7, cat. 100744 |
|  | PE/Dazzle™ 594 anti-mouse CD279 (PD-1) | 1/200, BioLegend, clone 29F.1A12, cat. 135228 |
|  | Alexa Fluor® 488 anti-mouse FOXP3 | 1/100, BioLegend, clone MF-14, cat 126406 |
|  | PerCP/Cyanine5.5 anti-mouse CD3ε | 1/200, BioLegend, clone 145-2C11, cat. 100328 |
|  | Brilliant Violet 421™ anti-mouse CD152 (CTLA4) | 1/200, BioLegend, clone UC10-4B9, cat. 106312 |
|  | eBioscience™ Fixable Viability Due eFluor™ 506 | 1/800, Thermo Fisher Scientific, cat. 65-0866-18 |

### H&E staining on PFA-fixed cryosections

Ten μm sections were cut from unfixed tissues in a frozen block; sections were stored at -20°C until ready to use. When ready, slides were taken out of the freezer at least 30 min before staining. Room temperature slides were fixed with cold 4% PFA for 10 min at room temperature followed by washing in PBS for 10 min at room temperature. After washing, slides were dipped into Mayer’s haematoxylin solution (see preparation under “Reagents and Solutions”) for 10 s, followed by washing in running tap water for 5 min. Then, slides were dipped into Eosin (5%) solution (see preparation under “Reagents and Solutions”) for 20 s, followed by washing in running tap water for 5 min. For rehydration, slides were placed first in 96% ethanol (2×2min) followed by 100% ethanol (2×2min). For clearance slides were placed in RotiClear® solution (Roth, cat. A538.5) for two-times 5 min, after which time, slides were mounted using Eukitt® quick-hardening mounting medium (Merck, cat. 03989). Slides were scanned using Leica DM6 B microscope and Leica Aperio Versa 8 slides scanner with 20x zoom and images were analysed using the ImageScope programme (version 12.3.3).

### Statistical analysis

All statistical analysis was carried out using One-Way ANOVA and Fisher LSD tests, using the Statistica programme (release 7).

## DATA AVAILABILITY

All data needed to evaluate the conclusions on the paper are presented in the paper and/or the Supplementary Information. Additional data related to the findings of this study are available from the corresponding authors.

## ACKNOWLEDGEMENTS

We would like to thank Stuart P. Atkinson for English editing, Merje Jakobson for performing H&E studies on paraffin-embedded formalin-fixed tissues, Dr. Aivar Orav for in vivo toxicity study analysis and Dr. Mario Plaas for the help with the slide scanner. PS acknowledges support from the Estonian Research Council (grant: PUT PSG38 to PS) and a Feasibility fund of the University of Tartu (grant: ARENG51 to PS). AL acknowledges a PhD fellowship from the Estonian government. MJV acknowledges the support by European Research Council grants (ERC-CoG-2014-648831 “My-Nano” and ERC-PoC-2018-825798 “Polymmune”). Part of the equipment employed in this work has been funded by Generalitat Valenciana and co-financed with FEDER funds (PO FEDER of Comunitat Valenciana 2014−2020). UH acknowledges the support by EsRC Mobilitas+ grant MOBTP185. TT acknowledges the support by UT EIK grant and GMVBS0230PR.

## AUTHOR CONTRIBUTIONS

AL performed the in vitro and in vivo experiments, histology, immunofluorescence, flow cytometry, analysis and wrote the manuscript. AM performed chemical design, synthesis, and characterisation and edited the manuscript. UH performed flow cytometry experiments and analysis. EA, MP, and MB performed computational simulations. SD performed drug release studies and chemical characterisation. LS performed analysis and expert evaluation of H&E images. PP edited the manuscript and provided discussions and lab support. TT edited the manuscript, participated in the experimental design and discussions, and provided laboratory support. MJV performed chemical design, in vitro and in vivo experiment design, supervised chemical synthesis and characterisation, provided lab support, and edited the manuscript. PS supervised all the experiments, participated in their design and analysis, and edited the manuscript. All authors edited the manuscript and approved the final version.

## CORRESPONDING AUTHORS

Correspondence to Tambet Teesalu, María J. Vicent and Pablo Scodeller.

## REFERENCES

1. Rivenbark, A. G., O’Connor, S. M. & Coleman, W. B. Molecular and cellular heterogeneity in breast cancer: challenges for personalized medicine. Am. J. Pathol. 183, 1113–1124 (2013).

2. Foulkes, W. D., Smith, I. E. & Reis-Filho, J. S. Triple-Negative Breast Cancer. N. Engl. J. Med. 363, 1938–1948 (2010).

3. Lehmann, B. D. et al. Identification of human triple-negative breast cancer subtypes and preclinical models for selection of targeted therapies. J. Clin. Invest. 121, 2750–2767 (2011).

4. Garrido-Castro, A. C., Lin, N. U. & Polyak, K. Insights into Molecular Classifications of Triple-Negative Breast Cancer: Improving Patient Selection for Treatment. Cancer Discov. 9, 176–198 (2019).

5. Pardoll, D. M. The blockade of immune checkpoints in cancer immunotherapy. Nat. Rev. Cancer 12, 252–264 (2012).

6. Esfahani, K. et al. A Review of Cancer Immunotherapy: From the Past, to the Present, to the Future. Curr. Oncol. 27, 87–97 (2020).

7. Adams, S. et al. Current Landscape of Immunotherapy in Breast Cancer: A Review. JAMA Oncol. 5, 1205–1214 (2019).

8. Gong, J., Chehrazi-Raffle, A., Reddi, S. & Salgia, R. Development of PD-1 and PD-L1 inhibitors as a form of cancer immunotherapy: a comprehensive review of registration trials and future considerations. J. Immunother. Cancer 6, 8 (2018).

9. Schmid, P. et al. Atezolizumab and Nab-Paclitaxel in Advanced Triple-Negative Breast Cancer. N. Engl. J. Med. 379, 2108–2121 (2018).

10. Schmid, P. et al. Atezolizumab plus nab-paclitaxel as first-line treatment for unresectable, locally advanced or metastatic triple-negative breast cancer (IMpassion130): updated efficacy results from a randomised, double-blind, placebo-controlled, phase 3 trial. Lancet Oncol. 21, 44–59 (2020).

11. Marra, A., Viale, G. & Curigliano, G. Recent advances in triple negative breast cancer: the immunotherapy era. BMC Med. 17, 90 (2019).

12. Mori, H. et al. The combination of PD-L1 expression and decreased tumor-infiltrating lymphocytes is associated with a poor prognosis in triple-negative breast cancer. Oncotarget 8, 15584–15592 (2017).

13. Mittendorf, E. A. et al. PD-L1 Expression in Triple-Negative Breast Cancer. Cancer Immunol. Res. 2, 361–370 (2014).

14. Socinski, M. A. et al. Atezolizumab for First-Line Treatment of Metastatic Nonsquamous NSCLC. N. Engl. J. Med. 378, 2288–2301 (2018).

15. Adams, S. et al. Patient-reported outcomes from the phase III IMpassion130 trial of atezolizumab plus nab-paclitaxel in metastatic triple-negative breast cancer. Ann. Oncol. 31, 582–589 (2020).

16. Fecher, L. A., Agarwala, S. S., Hodi, F. S. & Weber, J. S. Ipilimumab and Its Toxicities: A Multidisciplinary Approach. The Oncologist 18, 733–743 (2013).

17. Hunter, G., Voll, C. & Robinson, C. A. Autoimmune inflammatory myopathy after treatment with ipilimumab. Can. J. Neurol. Sci. J. Can. Sci. Neurol. 36, 518–520 (2009).

18. Maker, A. V. et al. Tumor Regression and Autoimmunity in Patients Treated With Cytotoxic T Lymphocyte–Associated Antigen 4 Blockade and Interleukin 2: A Phase I/II Study. Ann. Surg. Oncol. 12, 1005–1016 (2005).

19. Phan, G. Q. et al. Cancer regression and autoimmunity induced by cytotoxic T lymphocyte-associated antigen 4 blockade in patients with metastatic melanoma. Proc. Natl. Acad. Sci. U. S. A. 100, 8372–8377 (2003).

20. Cretella, D. et al. Pre-treatment with the CDK4/6 inhibitor palbociclib improves the efficacy of paclitaxel in TNBC cells. Sci. Rep. 9, 13014 (2019).

21. Arola, O. J. et al. Acute Doxorubicin Cardiotoxicity Involves Cardiomyocyte Apoptosis. Cancer Res. 60, 1789–1792 (2000).

22. Zhang, S. et al. Identification of the molecular basis of doxorubicin-induced cardiotoxicity. Nat. Med. 18, 1639–1642 (2012).

23. Keklikoglou, I. et al. Chemotherapy elicits pro-metastatic extracellular vesicles in breast cancer models. Nat. Cell Biol. 21, 190–202 (2019).

24. Hughes, R. et al. Perivascular M2 Macrophages Stimulate Tumor Relapse after Chemotherapy. Cancer Res. 75, 3479–3491 (2015).

25. Lewis, C. E. & Pollard, J. W. Distinct Role of Macrophages in Different Tumor Microenvironments. Cancer Res. 66, 605–612 (2006).

26. Peranzoni, E. et al. Macrophages impede CD8 T cells from reaching tumor cells and limit the efficacy of anti–PD-1 treatment. Proc. Natl. Acad. Sci. 115, E4041–E4050 (2018).

27. Neubert, N. J. et al. T cell–induced CSF1 promotes melanoma resistance to PD1 blockade. Sci. Transl. Med. 10, eaan3311 (2018).

28. Daurkin, I. et al. Tumor-Associated Macrophages Mediate Immunosuppression in the Renal Cancer Microenvironment by Activating the 15-Lipoxygenase-2 Pathway. Cancer Res. 71, 6400–6409 (2011).

29. Gok Yavuz, B. et al. Cancer associated fibroblasts sculpt tumour microenvironment by recruiting monocytes and inducing immunosuppressive PD-1+ TAMs. Sci. Rep. 9, 3172 (2019).

30. Pathria, P., Louis, T. L. & Varner, J. A. Targeting Tumor-Associated Macrophages in Cancer. Trends Immunol. 40, 310–327 (2019).

31. DeNardo, D. G. et al. Leukocyte Complexity Predicts Breast Cancer Survival and Functionally Regulates Response to Chemotherapy. Cancer Discov. 1, 54–67 (2011).

32. Mancini, V. S. B. W., Pasquini, J. M., Correale, J. D. & Pasquini, L. A. Microglial modulation through colony-stimulating factor-1 receptor inhibition attenuates demyelination. Glia 67, 291–308 (2019).

33. Lee, S., Shi, X. Q., Fan, A., West, B. & Zhang, J. Targeting macrophage and microglia activation with colony stimulating factor 1 receptor inhibitor is an effective strategy to treat injury-triggered neuropathic pain. Mol. Pain 14, 1744806918764979 (2018).

34. Bissinger, S. et al. Macrophage depletion induces edema through release of matrix-degrading proteases and proteoglycan deposition. Sci. Transl. Med. 13, eabd4550 (2021).

35. Wesolowski, R. et al. Phase Ib study of the combination of pexidartinib (PLX3397), a CSF-1R inhibitor, and paclitaxel in patients with advanced solid tumors. Ther. Adv. Med. Oncol. 11, 1758835919854238 (2019).

36. Papadopoulos, K. P. et al. First-in-Human Study of AMG 820, a Monoclonal Anti-Colony-Stimulating Factor 1 Receptor Antibody, in Patients with Advanced Solid Tumors. Clin. Cancer Res. 23, 5703–5710 (2017).

37. Kitamura, T. et al. Monocytes Differentiate to Immune Suppressive Precursors of Metastasis-Associated Macrophages in Mouse Models of Metastatic Breast Cancer. Front. Immunol. 8, (2018).

38. Madsen, D. H. et al. Tumor-Associated Macrophages Derived from Circulating Inflammatory Monocytes Degrade Collagen through Cellular Uptake. Cell Rep. 21, 3662– 3671 (2017).

39. Ishihara, D. et al. Wiskott-Aldrich Syndrome Protein Regulates Leukocyte-Dependent Breast Cancer Metastasis. Cell Rep. 4, 429–436 (2013).

40. Karousou, E. et al. Collagen VI and Hyaluronan: The Common Role in Breast Cancer. BioMed Res. Int. 2014, 1–10 (2014).

41. Scodeller, P. et al. Precision Targeting of Tumor Macrophages with a CD206 Binding Peptide. Sci. Rep. 7, 14655 (2017).

42. Lepland, A. et al. Targeting Pro-Tumoral Macrophages in Early Primary and Metastatic Breast Tumors with the CD206-Binding mUNO Peptide. Mol. Pharm. 17, 2518–2531 (2020).

43. Asciutto, E. K. et al. Phage-Display-Derived Peptide Binds to Human CD206 and Modeling Reveals a New Binding Site on the Receptor. J. Phys. Chem. B 123, 1973–1982 (2019).

44. Figueiredo, P. et al. Peptide-guided resiquimod-loaded lignin nanoparticles convert tumor-associated macrophages from M2 to M1 phenotype for enhanced chemotherapy. Acta Biomater. 133, 231–243 (2021).

45. Jameson, B. et al. Expression of DC-SIGN by Dendritic Cells of Intestinal and Genital Mucosae in Humans and Rhesus Macaques. J. Virol. 76, 1866–1875 (2002).

46. Conniot, J. et al. Immunization with mannosylated nanovaccines and inhibition of the immune-suppressing microenvironment sensitizes melanoma to immune checkpoint modulators. Nat. Nanotechnol. 14, 891–901 (2019).

47. Duro-Castano, A. et al. Well-Defined Star-Shaped Polyglutamates with Improved Pharmacokinetic Profiles As Excellent Candidates for Biomedical Applications. Mol. Pharm. 12, 3639–3649 (2015).

48. Arroyo-Crespo, J. J. et al. Tumor microenvironment-targeted poly-L-glutamic acid-based combination conjugate for enhanced triple negative breast cancer treatment. Biomaterials 186, 8–21 (2018).

49. Duro-Castano, A. et al. Polyglutamic acid-based crosslinked doxorubicin nanogels as an anti-metastatic treatment for triple negative breast cancer. J. Controlled Release 332, 10–20 (2021).

50. Arroyo-Crespo, J. J. et al. Anticancer Activity Driven by Drug Linker Modification in a Polyglutamic Acid-Based Combination-Drug Conjugate. Adv. Funct. Mater. 28, 1800931 (2018).

51. Shaffer, S. A. et al. In vitro and in vivo metabolism of paclitaxel poliglumex: identification of metabolites and active proteases. Cancer Chemother. Pharmacol. 59, 537–548 (2007).

52. Gordon, S. R. et al. PD-1 expression by tumour-associated macrophages inhibits phagocytosis and tumour immunity. Nature 545, 495–499 (2017).

53. Zhang, M. et al. Anti-CD47 Treatment Stimulates Phagocytosis of Glioblastoma by M1 and M2 Polarized Macrophages and Promotes M1 Polarized Macrophages In Vivo. PLOS ONE 11, e0153550 (2016).

54. Simon-Gracia, L. et al. Bifunctional Therapeutic Peptides for Targeting Malignant B Cells and Hepatocytes: Proof of Concept in Chronic Lymphocytic Leukemia. Adv. Ther. 3, 2000131 (2020).

55. MPD: Phenotype strain survey measures: alanine aminotransferase. https://phenome.jax.org/search/details/ssmeasures?searchterm=alanine+aminotransferase+&ontavail=2.

56. BALB/c Mouse | Charles River Laboratories. https://www.criver.com/products-services/find-model/balbc-mouse?region=3616.

57. Cassetta, L. & Kitamura, T. Targeting Tumor-Associated Macrophages as a Potential Strategy to Enhance the Response to Immune Checkpoint Inhibitors. Front. Cell Dev. Biol. 0, (2018).

58. Santoni, M. et al. Triple negative breast cancer: Key role of Tumor-Associated Macrophages in regulating the activity of anti-PD-1/PD-L1 agents. Biochim. Biophys. Acta BBA - Rev. Cancer 1869, 78–84 (2018).

59. Rodell, C. B. et al. TLR7/8-agonist-loaded nanoparticles promote the polarization of tumour-associated macrophages to enhance cancer immunotherapy. *Nat*. Biomed. Eng. 2, 578–588 (2018).

60. Loeuillard, E. et al. Targeting tumor-associated macrophages and granulocytic myeloid-derived suppressor cells augments PD-1 blockade in cholangiocarcinoma. J. Clin. Invest. 130, 5380–5396 (2020).

61. Choo, Y. W. et al. M1 Macrophage-Derived Nanovesicles Potentiate the Anticancer Efficacy of Immune Checkpoint Inhibitors. ACS Nano 12, 8977–8993 (2018).

62. Arlauckas, S. P. et al. Arg1 expression defines immunosuppressive subsets of tumor-associated macrophages. Theranostics 8, 5842–5854 (2018).

63. Landry, A. P., Balas, M., Alli, S., Spears, J. & Zador, Z. Distinct regional ontogeny and activation of tumor associated macrophages in human glioblastoma. Sci. Rep. 10, 19542 (2020).

64. Zheng, X. et al. Spatial Density and Distribution of Tumor-Associated Macrophages Predict Survival in Non–Small Cell Lung Carcinoma. Cancer Res. 80, 4414–4425 (2020).

65. Viitala, M. et al. Immunotherapeutic Blockade of Macrophage Clever-1 Reactivates the CD8 ^+^ T-cell Response against Immunosuppressive Tumors. Clin. Cancer Res. 25, 3289–3303 (2019).

66. Etzerodt, A. et al. Specific targeting of CD163+ TAMs mobilizes inflammatory monocytes and promotes T cell–mediated tumor regression. J. Exp. Med. 216, 2394–2411 (2019).

67. Linde, N. et al. Macrophages orchestrate breast cancer early dissemination and metastasis. Nat. Commun. 9, 1–14 (2018).

68. Witschen, P. M. et al. Tumor Cell Associated Hyaluronan-CD44 Signaling Promotes Pro-Tumor Inflammation in Breast Cancer. Cancers 12, 1325 (2020).

69. Guo, C. et al. Liposomal Nanoparticles Carrying anti-IL6R Antibody to the Tumour Microenvironment Inhibit Metastasis in Two Molecular Subtypes of Breast Cancer Mouse Models. Theranostics 7, 775–788 (2017).

70. Movahedi, K. et al. Nanobody-Based Targeting of the Macrophage Mannose Receptor for Effective *In Vivo* Imaging of Tumor-Associated Macrophages. Cancer Res. 72, 4165–4177 (2012).

71. Azad, A. K. et al. γ-Tilmanocept, a New Radiopharmaceutical Tracer for Cancer Sentinel Lymph Nodes, Binds to the Mannose Receptor (CD206). J. Immunol. 195, 2019–2029 (2015).

72. Jaynes, J. M., Lopez, H. W., Martin, G. R., Yates, C. & Garvin, C. E. Peptides having anti-inflammatory properties. (2016).

73. Scodeller, P. & Asciutto, E. K. Targeting Tumors Using Peptides. Molecules 25, 808 (2020).

74. Ekladious, I., Colson, Y. L. & Grinstaff, M. W. Polymer–drug conjugate therapeutics: advances, insights and prospects. Nat. Rev. Drug Discov. 18, 273–294 (2019).

75. Duro-Castano, A., Conejos-Sánchez, I. & Vicent, M. J. Peptide-Based Polymer Therapeutics. Polymers 6, 515–551 (2014).

76. Moura, L. I. F. et al. Functionalized branched polymers: promising immunomodulatory tools for the treatment of cancer and immune disorders. Mater. Horiz. 6, 1956–1973 (2019).

77. Melnyk, T., Đorđević, S., Conejos-Sánchez, I. & Vicent, M. J. Therapeutic potential of polypeptide-based conjugates: Rational design and analytical tools that can boost clinical translation. Adv. Drug Deliv. Rev. 160, 136–169 (2020).

78. Duro-Castano, A. et al. Capturing “Extraordinary” Soft-Assembled Charge-Like Polypeptides as a Strategy for Nanocarrier Design. Adv. Mater. 29, 1702888 (2017).

79. Duro-Castano, A., Movellan, J. & Vicent, M. J. Smart branched polymer drug conjugates as nano-sized drug delivery systems. Biomater. Sci. 3, 1321–1334 (2015).

80. Cortez-Retamozo, V. et al. Origins of tumor-associated macrophages and neutrophils. Proc. Natl. Acad. Sci. 109, 2491–2496 (2012).

81. Kurashige, M. et al. Origin of cancer-associated fibroblasts and tumor-associated macrophages in humans after sex-mismatched bone marrow transplantation. *Commun*. Biol. 1, 1–13 (2018).

82. Veglia, F. & Gabrilovich, D. I. Dendritic cells in cancer: the role revisited. Curr. Opin. Immunol. 45, 43–51 (2017).

83. Agostinis, P. et al. PHOTODYNAMIC THERAPY OF CANCER: AN UPDATE. CA. Cancer J. Clin. 61, 250–281 (2011).

84. Cheah, H. Y. et al. Near-Infrared Activatable Phthalocyanine–Poly-L-Glutamic Acid Conjugate: Enhanced in Vivo Safety and Antitumor Efficacy toward an Effective Photodynamic Cancer Therapy. Mol. Pharm. 15, 2594–2605 (2018).

85. Nguyen, V.-N., Yan, Y., Zhao, J. & Yoon, J. Heavy-Atom-Free Photosensitizers: From Molecular Design to Applications in the Photodynamic Therapy of Cancer. Acc. Chem. Res. 54, 207–220 (2021).

86. Tacar, O., Sriamornsak, P. & Dass, C. R. Doxorubicin: an update on anticancer molecular action, toxicity and novel drug delivery systems. J. Pharm. Pharmacol. 65, 157–170 (2013).

87. Shan, H., Dou, W., Zhang, Y. & Qi, M. Targeted ferritin nanoparticle encapsulating CpG oligodeoxynucleotides induces tumor-associated macrophage M2 phenotype polarization into M1 phenotype and inhibits tumor growth. Nanoscale 12, 22268–22280 (2020).

88. Ramesh, A., Brouillard, A., Kumar, S., Nandi, D. & Kulkarni, A. Dual inhibition of CSF1R and MAPK pathways using supramolecular nanoparticles enhances macrophage immunotherapy. Biomaterials 227, 119559 (2020).

89. Hollmén, M. et al. G-CSF regulates macrophage phenotype and associates with poor overall survival in human triple-negative breast cancer. OncoImmunology 5, e1115177 (2016).

90. Georgoudaki, A.-M. et al. Reprogramming Tumor-Associated Macrophages by Antibody Targeting Inhibits Cancer Progression and Metastasis. Cell Rep. 15, 2000–2011 (2016).

91. Zhang, F., et al. Reprogramming of profibrotic macrophages for treatment of bleomycin-induced pulmonary fibrosis. EMBO Mol. Med. 12, e12034 (2020).

92. Sartor, O. et al. Lutetium-177–PSMA-617 for Metastatic Castration-Resistant Prostate Cancer. N. Engl. J. Med. 0, null (2021).

93. Tian, C. et al. ff19SB: Amino-Acid-Specific Protein Backbone Parameters Trained against Quantum Mechanics Energy Surfaces in Solution. J. Chem. Theory Comput. 16, 528–552 (2020).

94. AmberTools - SBGrid Consortium - Supported Software. https://sbgrid.org/software/titles/ambertools.

95. Hopkins, C. W., Le Grand, S., Walker, R. C. & Roitberg, A. E. Long-Time-Step Molecular Dynamics through Hydrogen Mass Repartitioning. J. Chem. Theory Comput. 11, 1864–1874 (2015).

96. Berendsen, H. J. C., Postma, J. P. M., van Gunsteren, W. F., DiNola, A. & Haak, J. R. Molecular dynamics with coupling to an external bath. J. Chem. Phys. 81, 3684–3690 (1984).

97. Ryckaert, J.-P., Ciccotti, G. & Berendsen, H. J. C. Numerical integration of the cartesian equations of motion of a system with constraints: molecular dynamics of n-alkanes. J. Comput. Phys. 23, 327–341 (1977).

98. Ray, A. & Dittel, B. N. Isolation of Mouse Peritoneal Cavity Cells. J. Vis. Exp. JoVE 1488 (2010) doi:10.3791/1488.

99. Bankhead, P. et al. QuPath: Open source software for digital pathology image analysis. Sci. Rep. 7, 16878 (2017).

