## Supplementary Information for "Depletion of CD206^+^ Tumour Macrophages via a Peptide-Targeted Star-Shaped Polyglutamate Inhibits Tumourigenesis and Metastatic Dissemination in Breast Cancer Models"

¥: equal contribution
ψ: equal contribution

#### Synthetic procedures

##### Materials

Three arms benzenetricarboxamide (BTA) centred star-shaped poly-L-glutamates (St-PGA) was kindly provided by Polypeptide Therapeutic Solutions S.L. The mUNO peptide (CSPGAK-COOH) was purchased from TAG Copenhagen. The Oregon Green™ 488 (OG) Cadaverine and trifluoroacetic acid were purchased from Thermo Fisher Scientific. Doxorubicin hydrochloride salt (DOX) was purchased from Xingcheng Chempharm Co. Ltd. Daunorubicin HCl (DAU), cathepsin B from bovine spleen, dithiothreitol, and sodium acetate were purchased from Sigma-Aldrich and used without further purification unless otherwise indicated. Deuterium oxide was purchased from Deutero GmbH. Size exclusion chromatography (SEC) was performed using Sephadex® LH-20, and columns were purchased from GE Healthcare Bio-Sciences AB. Dialysis was performed in a Millipore ultrafiltration device fitted with a 3 kDa molecular weight cut-off (MWCO) regenerated cellulose membrane (Vivaspin®, Merck). For the drug release study, $EDTANa_2$, PBS, chloroform, liquid-chromatography–mass spectrometry (LC-MS) grade methanol, and water were purchased from Merck, while LC-MS grade acetonitrile was purchased from Fisher chemical.

#### Synthesis of St-PGA-OG-mUNO

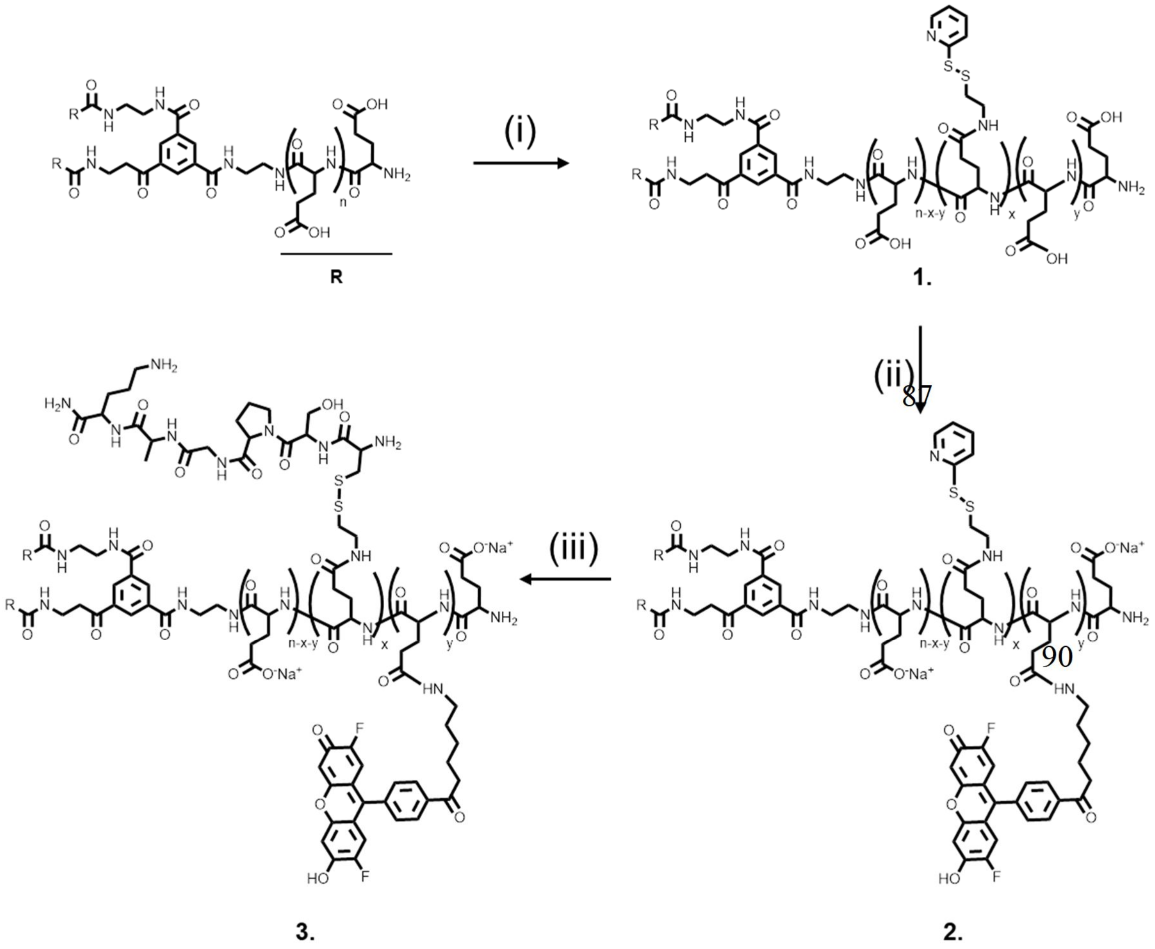


**Scheme S1. Synthetic approach for St-PGA-OG-mUNO. (i)** *a.* DMTMM $BF_4$, 30 min, RT, anh-DMF, *b.* pyridyldithiol (PD) amine, DIEA, pH 8, anh-DMF, 24 h; **(ii)** *a.* DMTMM $BF_4$, 30, RT, anh-DMF, *b.* $OG_{488}$-cadaverine, DIEA, pH 8, anh-DMF, 24 h; **(iii)** mUNO, 3 h, PBS, pH 7.4.

#### Synthesis of St-PGA-PD

The synthesis of St-PGA-PD (PD – pyridyldithiol) was performed according to previously published protocols[1,2]. St-PGA (50 mg, 0.387 mmol glutamic acid units (GAU), 1 eq.) was dissolved in 5 mL of anhydrous N,N'-dimethylformamide (DMF) under nitrogen flow. Then, 4-(4,6-Dimethoxy-1,3,5-triazin-2-yl)-4-methylmorpholinium tetrafluoroborate (DMTMM $BF_4$, 9.06 mg, 0.075 eq.) was added after dissolving in anhydrous DMF. The solution was stirred for 30 min at room temperature. Then, pyridyldithiol amine (4.1 mg, 0.06 eq.) was added to the solution and the pH adjusted to 8 by adding N,N-diisopropylethylamine (DIEA). The mixture was kept under magnetic stirring for 24 h at room temperature. The mixture was purified by precipitation in diethyl ether (3x100 mL). The compound was further purified through either acid/base precipitation. A white amorphous solid was obtained after freeze-drying.

Yield: 80-90%.

1H NMR δH (300MHz, $D_2O$): 8.53-8.19 (1H, b), 7.93-7.74 (2H, b), 7.45-7.23 (1H, b), 4.48-4.23 (1H, b), 2.40-1.79 (4H, m).

**Synthesis of St-PGA-PD-OG**

The synthesis of St-PGA-PD-OG was performed according to a previously published protocol[2]. St-PGA-PD (25 mg, 0.1935 mmol GAU, 1 eq.) was dissolved in 2.5 mL of anhydrous DMF under nitrogen flow. Then, DMTMM $BF_4$ (4.53 mg, 0.075 eq.) dissolved in anhydrous DMF was added. The solution was stirred for 30 min at room temperature. Then, $OG_{488}$-cadaverine (1.1 mg, 0.012 eq.) was added to the solution, and the pH was adjusted to 8 by adding DIEA. The mixture was kept under magnetic stirring for 24 h at room temperature and protected from the light. The mixture was purified by precipitation in diethyl ether (3x100 mL). The fine powder obtained was dissolved in DMF and passed twice through an LH-20 column. The first eluting fraction, corresponding to the acid form of the St-PGA-PD-OG, was collected and dried under vacuum, and the water-soluble sodium salt form of the final product was obtained by dissolving the resulting solid in 0.1 M $NaHCO_3$. The excess of the free drug was further removed by using Vivaspin® 3 kDa. An orange-red amorphous solid was obtained after freeze-drying.

Yield: 75-80% wt.

**Synthesis St-PGA-OG-mUNO**

The synthesis of St-PGA-OG-mUNO was performed using a previously described protocol[2]. Briefly, St-PGA-PD-OG (20 mg, 1.0 eq) was dissolved at the concentration of 10 mg/mL in PBS, pH 7.4, room temperature under gently stirring. The mUNO peptide (4.1 mg, 0.055 eq.) was dissolved in PBS, immediately added to the solution, and stirred for 3 h in the dark. Then, the reaction was purified by Vivaspin® 3 kDa. The final product was lyophilised, and an amorphous orange-red solid was obtained.

Yield: 60% wt.

**Synthesis of St-PGA-DOX-mUNO ("OximUNO")**

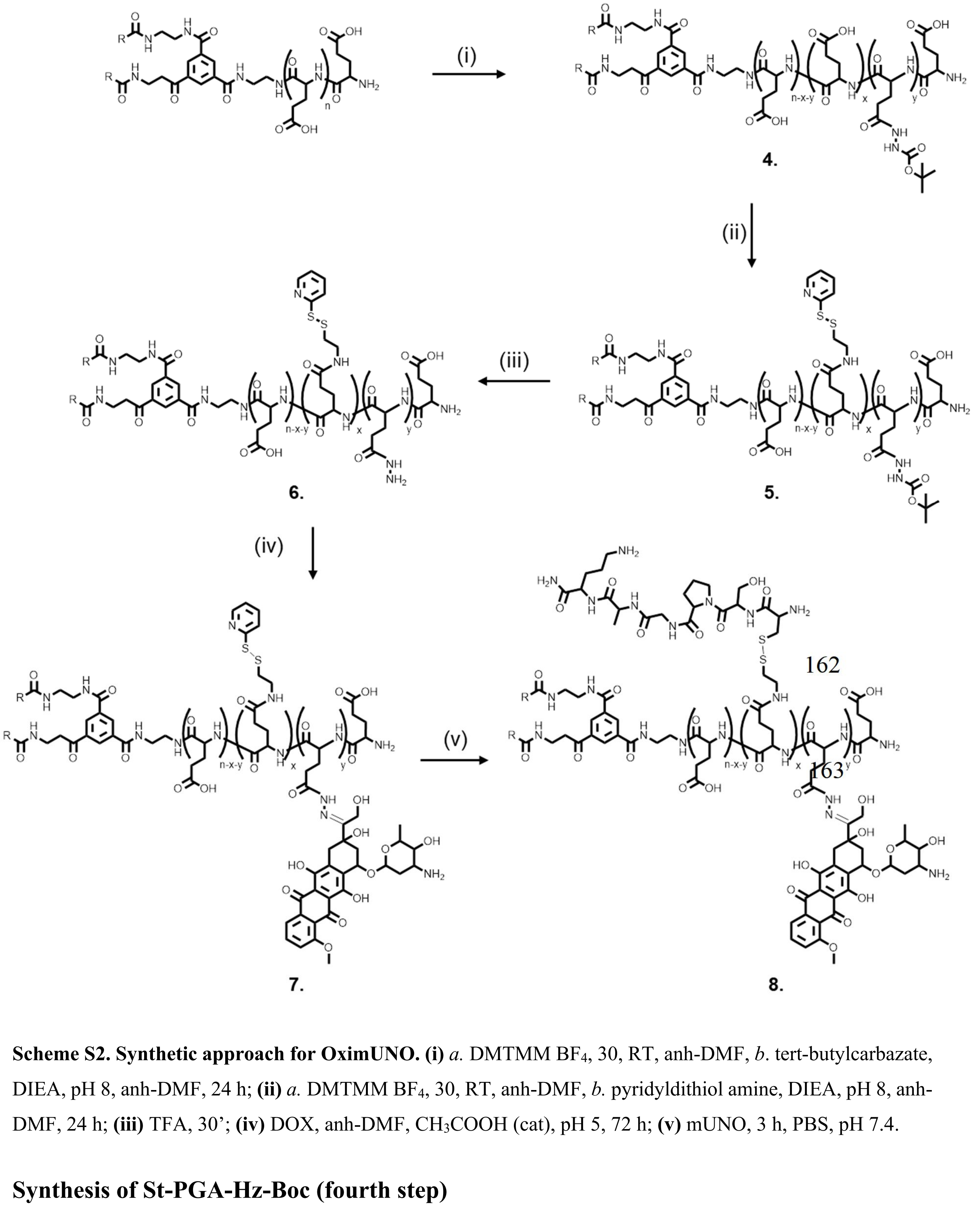


**Scheme S2. Synthetic approach for OximUNO. (i)** *a.* DMTMM $BF_4$, 30, RT, anh-DMF, *b.* tert-butylcarbazate, DIEA, pH 8, anh-DMF, 24 h; **(ii)** *a.* DMTMM $BF_4$, 30, RT, anh-DMF, *b.* pyridyldithiol amine, DIEA, pH 8, anh-DMF, 24 h; **(iii)** TFA, 30'; **(iv)** DOX, anh-DMF, $CH_3COOH$ (cat), pH 5, 72 h; **(v)** mUNO, 3 h, PBS, pH 7.4.

**Synthesis of St-PGA-Hz-Boc (fourth step)**

The synthesis of St-PGA-Hz-Boc was performed using a modified previously published protocol[3]. Briefly, St-PGA (500 mg, 3.67 mol GAU, 1 eq.) was dissolved in 10 mL of anhydrous DMF under nitrogen flow. Afterwards, DMTMM $BF_4$ (90.56 mg, 0.075 eq.) dissolved in anhydrous DMF was added. The solution was stirred for 30 min at room temperature. Then, tert-butylcarbazate (29.15 mg, 0.06 eq.) was added to the solution, and the pH was adjusted to 8 by adding DIEA. The mixture was kept under magnetic stirring for 24 h at room temperature. The mixture was purified by precipitation in diethyl ether (3x300 mL). The compound was further purified through either acid/base precipitation. A white amorphous solid was obtained after freeze-drying.

Yield: 80-90% wt.

1H NMR δH (300MHz, $D_2O$): 4.36-4.14 (1H, b), 2.40-1.79 (4H, m), 1.42-1.37 (9H, s).

**Synthesis of St-PGA- Hz-Boc-PD (fifth step)**

St-PGA (250 mg, 1.935 mol GAU, 1 eq.) was dissolved in 10 mL of anhydrous DMF under nitrogen flow. Then, DMTMM $BF_4$ (45.28 mg, 0.075 eq.) dissolved in anhydrous DMF was added. The solution was stirred for 30 min at room temperature. Then, pyridyldithiol amine (20.46 mg, 0.06 eq.) was added to the solution and the pH was adjusted to 8 by adding DIEA. The mixture was kept under magnetic stirring for 24 h at room temperature. The mixture was purified by precipitation in diethyl ether (3x300 mL). The compound was further purified through either acid/base precipitation. A white amorphous solid was obtained after freeze-drying.

Yield: 80-90% wt.

1H NMR δH (300MHz, $D_2O$): 8.49-8.28 (1H, b), 7.95-7.81 (2H, b), 7.45-7.28 (1H, b), 4.49-4.19 (1H, b), 2.41-1.81 (4H, m), 1.57-1.42 (9H, s).

**Deprotection of St-PGA-Hz and St-PGA- Hz-PD (sixth step)**

The deprotection of protected St-PGA-Hz-Boc and St-PGA -Hz-Boc-PD (200 mg each) was performed by dissolving the conjugates in trifluoroacetic acid (100%, 5 mL). After complete dissolution, the reaction was allowed to continue for 30 minutes. The nanoconjugates were precipitated in diethyl ether (3x300 mL), washed with acidic water, and lyophilised.

Yield: 70% wt.

1H NMR δH (300MHz, $D_2O$): 8.51-8.27 (2H, b), 7.90-7.79 (1H, b), 7.40-7.27 (2H, b), 4.50-4.22 (1H, b), 2.47-1.91 (4H, m).

**Synthesis of St-PGA-DOX and St-PGA- DOX-PD (seventh step)**

Deprotected polymers were conjugated with DOX following a previously published protocol[3]. Briefly, St-PGA-Hz or St-PGA-Hz-PD (100 mg, 0.734 mmol, 1 eq.) and DOX (42.57 mg, 0.1 eq.) were dissolved in 5 mL of anhydrous DMF at room temperature under magnetic stirring.

After the full solubilisation of reagents, acetic acid (glacial, 100 µL) was added to the solution to reach pH 5. The reaction was allowed to proceed for 72 h at room temperature, protected from light. After that time, the reaction volume was reduced by half, and the mixture was purified by passing it through an LH-20 column two times to remove unreacted DOX. The first eluting fraction, corresponding to the acid form of the DOX-conjugate, was collected, dried under vacuum, and the water-soluble sodium salt form of the final product was obtained by dissolving the resulting solid in 0.1 M $NaHCO_3$. The excess of the free drug was further removed using Vivaspin® 3 kDa. The final product was lyophilised, and an amorphous dark red solid was obtained.

Yield(s):

- St-PGA-DOX: 70-80 % wt
- St-PGA-DOX-PD: 70 % wt

**Synthesis of OximUNO (eighth step)**

St-PGA-PD-DOX (80 mg, 1.0 eq) was dissolved at the concentration of 10 mg/mL in PBS, pH 7.4, room temperature under gentle stirring. The mUNO peptide (16.35 mg, 0.055 eq.) was dissolved in PBS, immediately added to the solution, and stirred for 3 h in the dark. Then, the product was purified by Vivaspin® 3 kDa. The final product was lyophilised, and an amorphous dark red solid was obtained.

Yield: 60% wt.

**OG/DOX/mUNO loading determination**

OG loading was determined by fluorescence spectroscopy (λex= 501 nm, λem= 526 nm) using a calibration curve previously constructed with the OG standard solutions in the concentration range 0.5-20 µg/mL ($y=6\cdot10^6x - 7240$, $R^2= 0.9988$).

DOX loading was determined by UV-Vis at 480 nm using a calibration curve previously made with DOX standard solutions in the concentration range of 5.0-50 µg/mL ($y=0.0197x + 0.0045$, $R^2= 0.9988$). mUNO loading was estimated by $^1$H-NMR and further confirmed by LC-MS amino acid analysis performed at the University of Barcelona (Unitat de Tècniques Separatives I Síntesi de Pèptids Centres Cientifics I Tecnològics).

**Free drug determination**

OximUNO nanoconjugate (3 mg/mL) was suspended in 500 µL of methanol (LC-MS grade) with DAU as internal standard, vortexed for 2 min and centrifuged at 30,437g for 10 min. The supernatant was filtered through a 0.45 µm filter and subjected to LC-MS analysis.

#### Supplementary figures and video

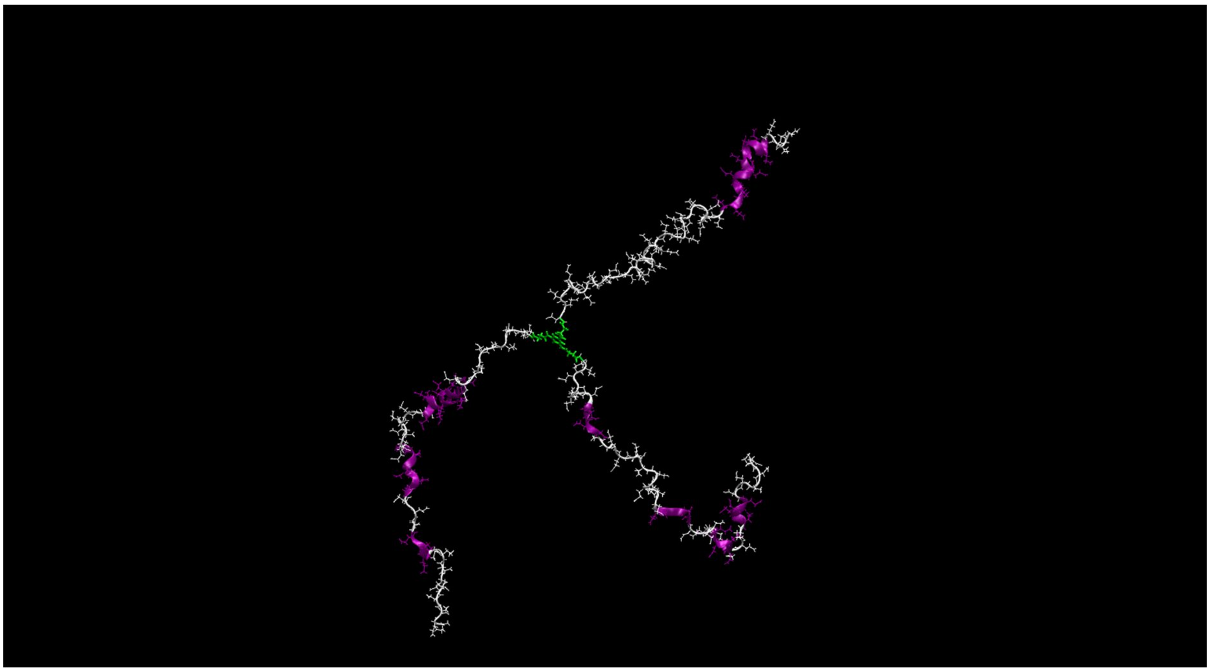

**Video S1. Fifty ns MD trajectory of St-PGA in solution.** Water and ions were removed for visualisation purposes. PGA chains are shown in white by overlaying in magenta the regions that form the alpha-helix structure, and the BTA core is shown in green.

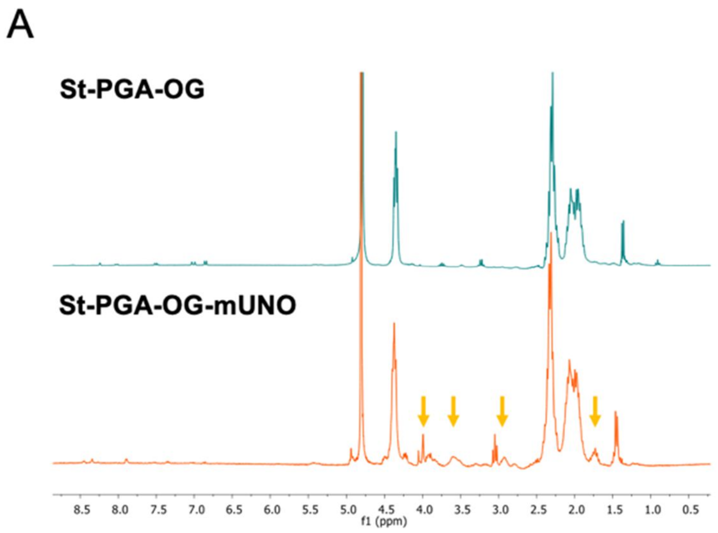


**Fig. S1. Representative characterisation of St-PGA-OG and St-PGA-OG-mUNO.** $^1$H-NMR in $D_2O$ of the conjugates. The orange arrows show the peaks from mUNO.

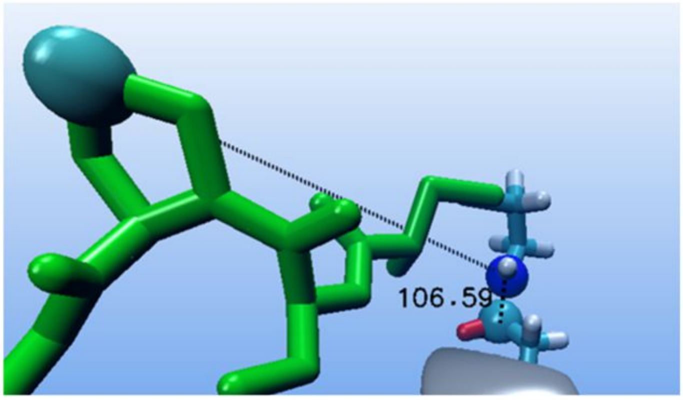


**Fig. S2. The angle used to characterise the rotation of mUNO around PGA.** The angle formed by an aromatic carbon of mUNO's proline (green sphere), a nitrogen of the pyridyldithiol linker (blue sphere) and an aromatic carbon of the glutamic acid (light blue sphere).

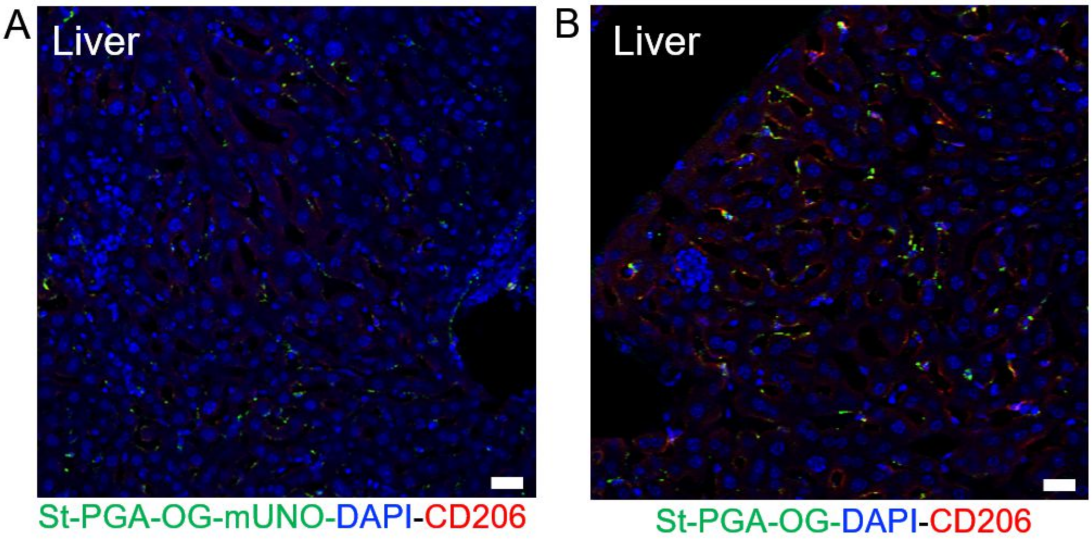


**Fig. S3. St-PGA-OG-mUNO shows low hepatic accumulation in the orthotopic TNBC model.** St-PGA-OG-mUNO (0.41 mg/0.5mL) or St-PGA-OG (0.35 mg/mL) was i.p. injected ten days post tumour induction (s.c. injection of 1x10$^{6}$ 4T1 cells), N=3. Nanoconjugates were circulated for 6 h, after which time, mice were sacrificed, and organs collected for analysis. Both St-PGA-OG-mUNO **(A)** and St-PGA-OG **(B)** showed low hepatic accumulation. Scale bars = 20 µm.

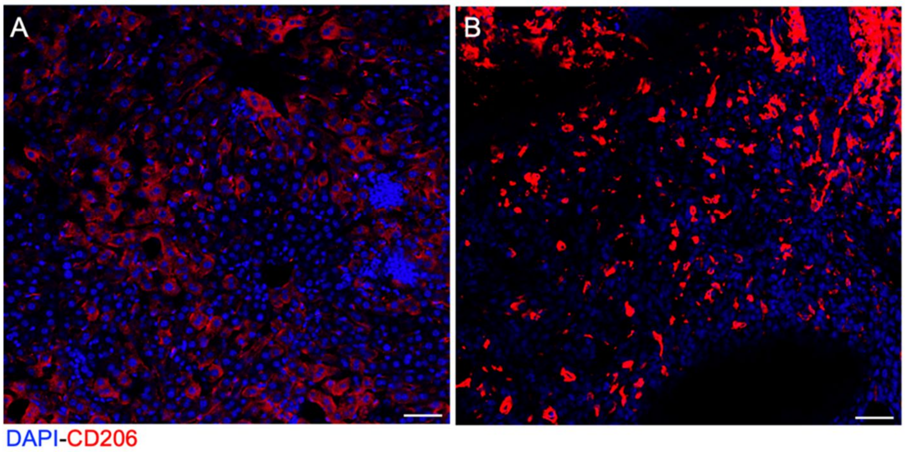


**Fig. S4**. **Immunostaining showing CD206 expression in the liver compared to the saturated signal of CD206 in the orthotopic 4T1 tumour.** Representative image of the liver **(A)** and tumour **(B)** from 4T1 s.c. tumour-bearing mice shown with higher intensity in the red channel (corresponding to CD206). Scale bars = 50 µm.

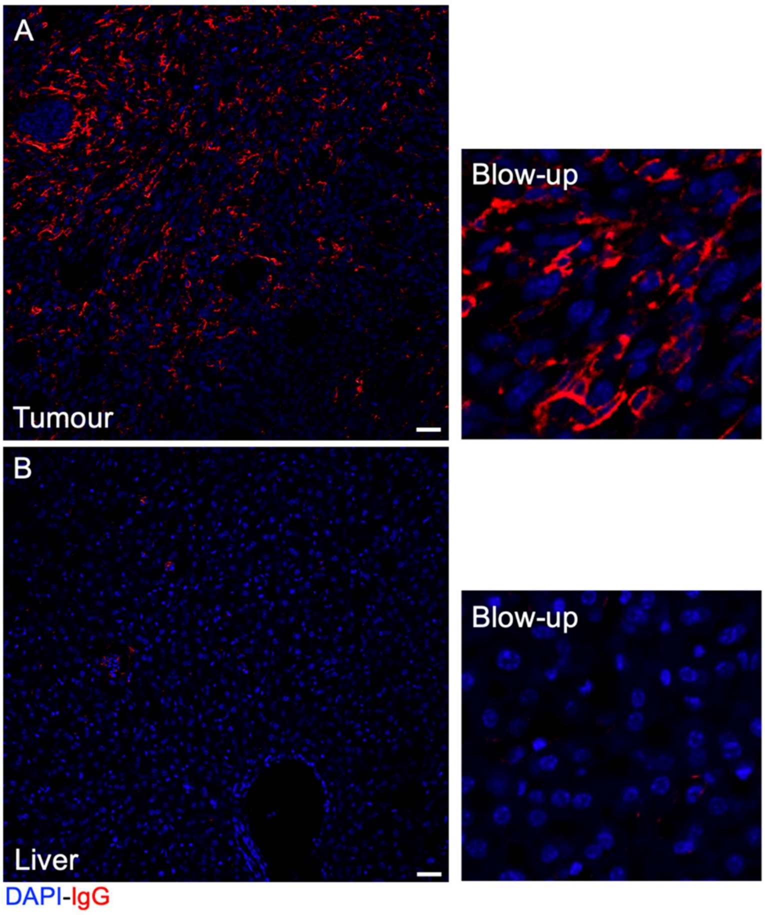


**Fig. S5**. **Immunostaining of endogenous IgG indicates leaky tumour vasculature in the orthotopic 4T1 tumours.** Tumour **(A)** and liver **(B)** sections of orthotopic 4T1 tumour-bearing mice were stained using rat anti-mouse IgG and counterstained with DAPI. Representative images of N=3 mice. Scale bars = 20 µm.

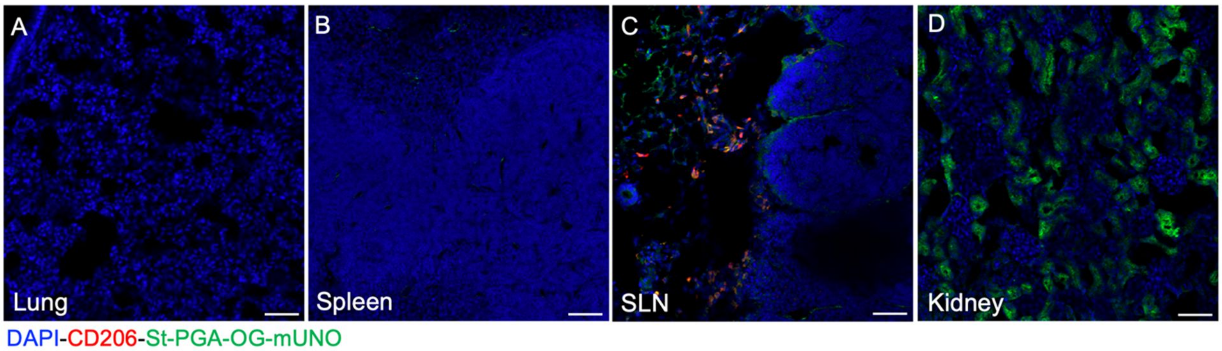


**Fig. S6. Distribution of St-PGA-OG-mUNO in the lung, spleen, sentinel lymph node (SLN) and kidney.** St-PGA-OG-mUNO (0.41 mg/0.5mL) was i.p. injected ten days post tumour induction (s.c. injection of $1x10^6$ 4T1 cells). Nanoconjugates were circulated for 6 h, after which time, mice were sacrificed, and organs collected for analysis (stained for CD206 and OG). Representative images of (A) lung, (B) spleen, (C) SLN, and (D) kidney showing distribution of St-PGA-OG-mUNO. Representative images from N=3 mice. Scale bars = 50 µm.

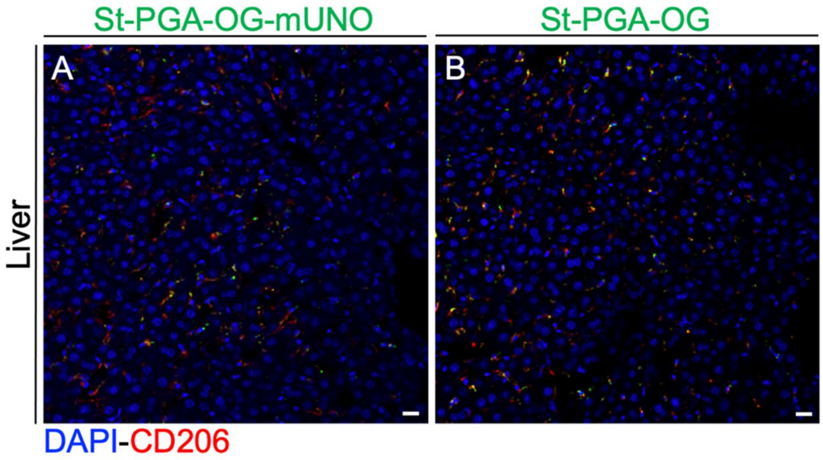


**Fig. S7. St-PGA-OG-mUNO shows low hepatic accumulation in the experimental metastasis of TNBC model.** St-PGA-OG-mUNO (0.41 mg/0.5mL) or St-PGA-OG (0.35 mg/0.5mL) were i.p. injected ten days post tumour induction in lungs (i.v. injection of $5x10^5$ 4T1 cells), N=2. Nanoconjugates were circulated for 6 h, after which time, mice were sacrificed, and organs collected for analysis. Both (**A**) St-PGA-OG-mUNO and (**B**) St-PGA-OG displayed low hepatic accumulation. Scale bars = 20 µm.

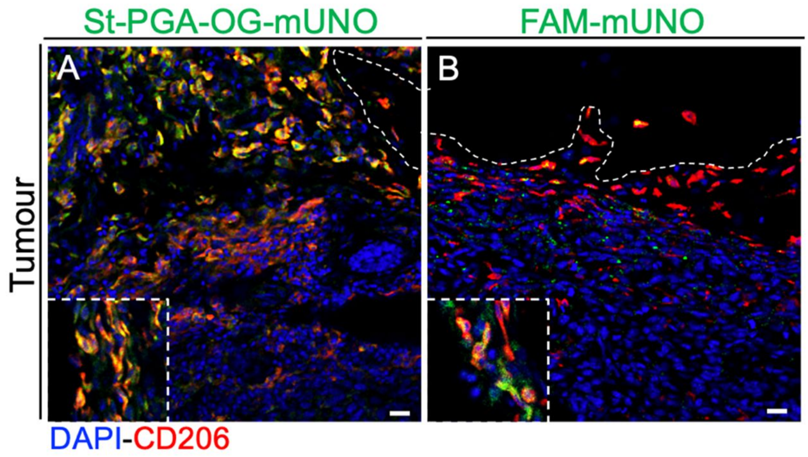


**Fig. S8. St-PGA-OG-mUNO shows higher receptor colocalisation than FAM-mUNO.** St-PGA-OG-mUNO (30 nmoles OG) or FAM-mUNO (30 nmoles FAM) were i.p. injected ten days post tumour induction (s.c. injection of $1x10^6$ 4T1 cells), N=2. The nanoconjugate or free peptide was circulated for 6 h, after which time, mice were sacrificed, and organs collected for analysis. (**A**) St-PGA-OG-mUNO showed higher receptor colocalisation than **(B)** FAM-mUNO. Scale bars = 20 µm.

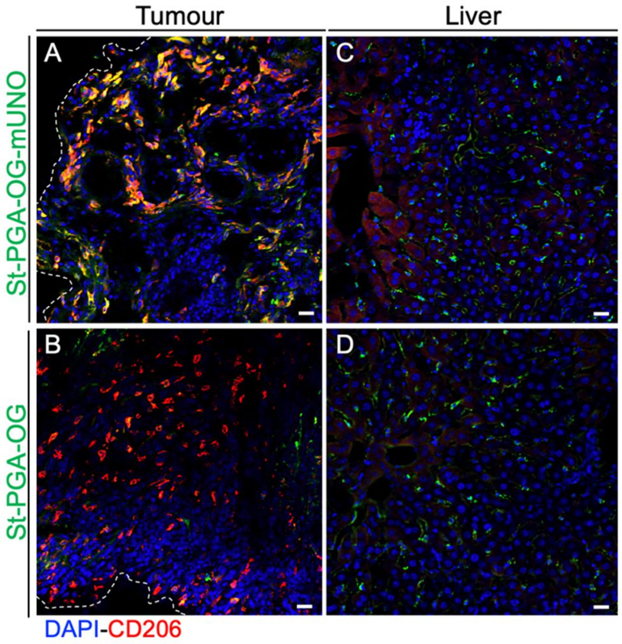


**Fig. S9. St-PGA-OG-mUNO shows high homing to M2 TAMs on the orthotopic TNBC model but higher hepatic accumulation with a higher dose.** St-PGA-OG-mUNO (0.82mg/0.5mL) or St-PGA-OG (0.7mg/0.5mL) were i.p. injected ten days post tumour induction (s.c. injection of 1x$10^6$ 4T1 cells), N=2. Nanoconjugates were circulated for 6 h after which time, mice were sacrificed, and organs collected for analysis. (**A**) St-PGA-OG-mUNO showed high colocalisation with CD206 whereas (**B**) St-PGA-OG showed minimal colocalisation. (**C, D**) Both nanoconjugates displayed moderate hepatic accumulation. Scale bars = 20 µm.

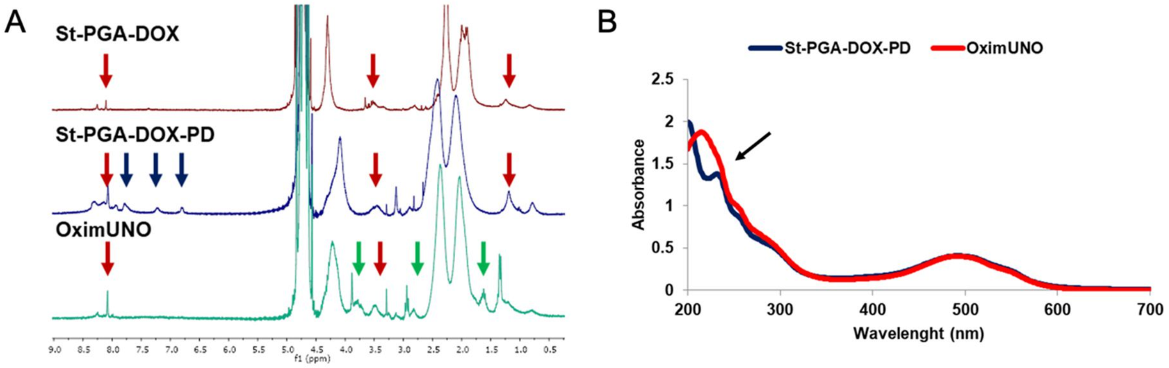


**Fig. S10. Representative characterisation of St-PGA-DOX and OximUNO.** (**A**) $^{1}$H-NMR in $D_2O$ of nanoconjugates. Red arrows indicate the signals from DOX, blue arrows show the peaks from the pyridyl of the pyridyldithiol (PD), while green arrows indicate signals from mUNO. (**B**) UV-Vis spectrum of St-PGA-PD-DOX (dark blue line) and OximUNO (red line) showing the displacement of pyridyl moiety at 260 nm by mUNO.

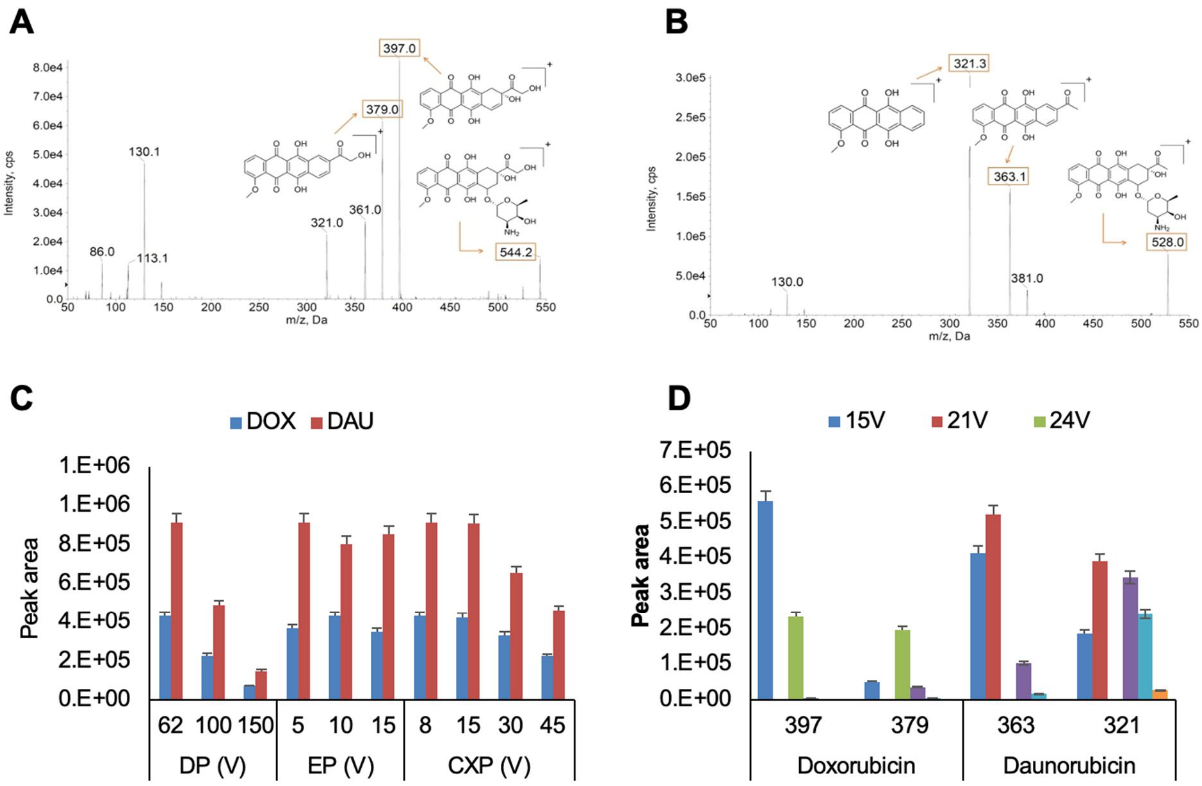


E

| MS parameter | Value |
|---|---|
| CXP | 8 V |
| DP | 62 V |
| EP | 5 V |
| CE (DAU) | 21 V |
| CE (DOX-397 m/z) | 15 V |
| CE (DOX-379 m/z) | 24 V |

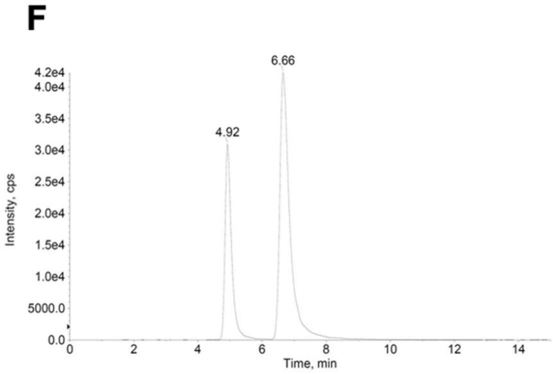


G

| | Recovery | | Matrix effects | | LOD (µg/ml) | LOQ (µg/ml) | Accuracy (%) |
|---|---|---|---|---|---|---|---|
| | DOX | DAU | DOX | DAU | DOX | DOX | DOX |
| **dPBS 7.4** | 94.44% | 68.25% | 1.03 | 0.95 | | | |
| **acetate buffer pH 5** | 128.50% | 67.66% | 1.41 | 0.96 | 4.62 | 10.72 | 78-120 |
| **i.p. fluid** | N/A | N/A | 1.17 | 1.76 | | | |

**Fig. S11. LC-MS method development for the determination of DOX in drug release studies and stability studies of OximUNO in i.p. fluid and dPBS** (**A**) MS-MS fragmentation spectra of DOX. (**B**) MS-MS fragmentation spectra of DAU. (**C**) Optimisation of MS parameter (average ± Sd, N=3). (**D**) Optimisation of CE for each mass transition of DOX and DAU (average ± Sd, N=3). (**E**) Final MS parameters. (**F**) Representative LC-MS chromatogram of DOX and DAU with $t_r$=4.92 min and $t_r$=6.66 min, respectively. (**G**) Method validation parameters. Error bars represent SE. Abbreviations: i.p. fluid – intraperitoneal fluid; DAU – daunorubicin; CXP

– collision exit potential; DP – declustering potential; EP – entrance potential; CE – collision energy; LOD – limit of detection; LOQ – limit of quantification; N/A – not applicable.

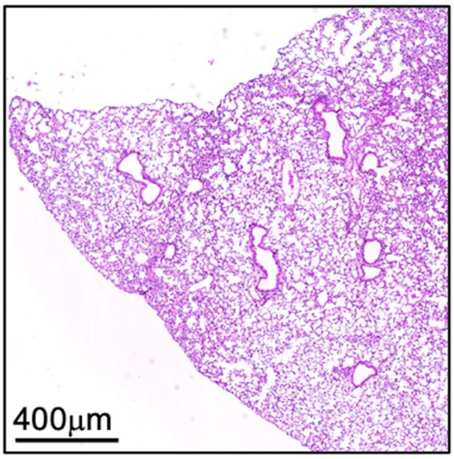


**Fig. S12. H&E on healthy Balb/c mouse lung.** Representative microscopy image from healthy female Balb/c mouse lung used as a control in H&E analysis showing the typical lung structure. Scale bar = 400 μm.

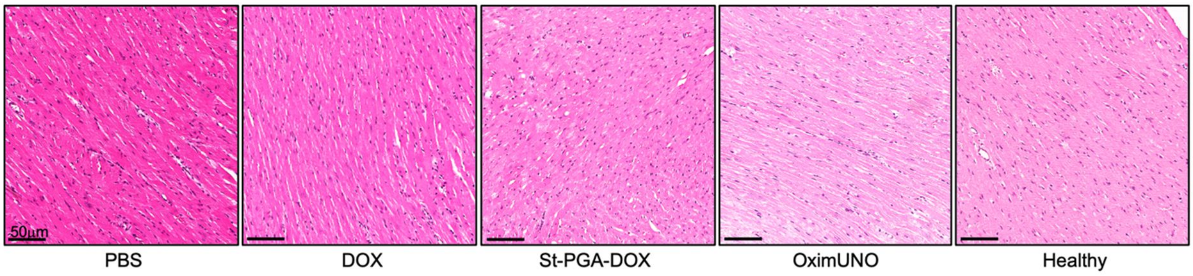


**Fig. S13. H&E staining of hearts from the treatment of orthotopic TNBC.** Hearts from all treatment groups were analysed with H&E for potential cardiotoxicity. All treatment groups showed a lack of cardiotoxicity when analysed with H&E. Heart tissue structure in mice bearing orthotopic TNBC tumours remains similar to that of the healthy heart (farthest right). Scale bar = 50 μm.

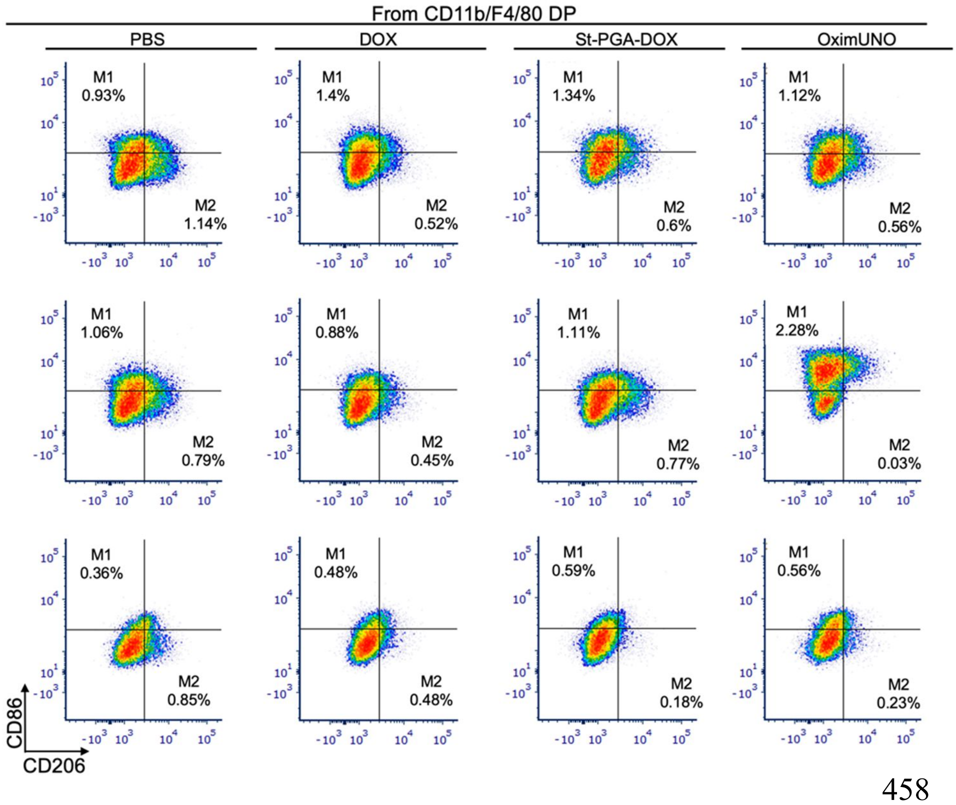


**Fig. S14. Flow cytometry plots for M2 and M1 macrophages.** Cytometry plots for the M2 and M1 macrophage populations of the treatment study shown in Figure 5. Percentages shown are from total cells. DP: double positive.

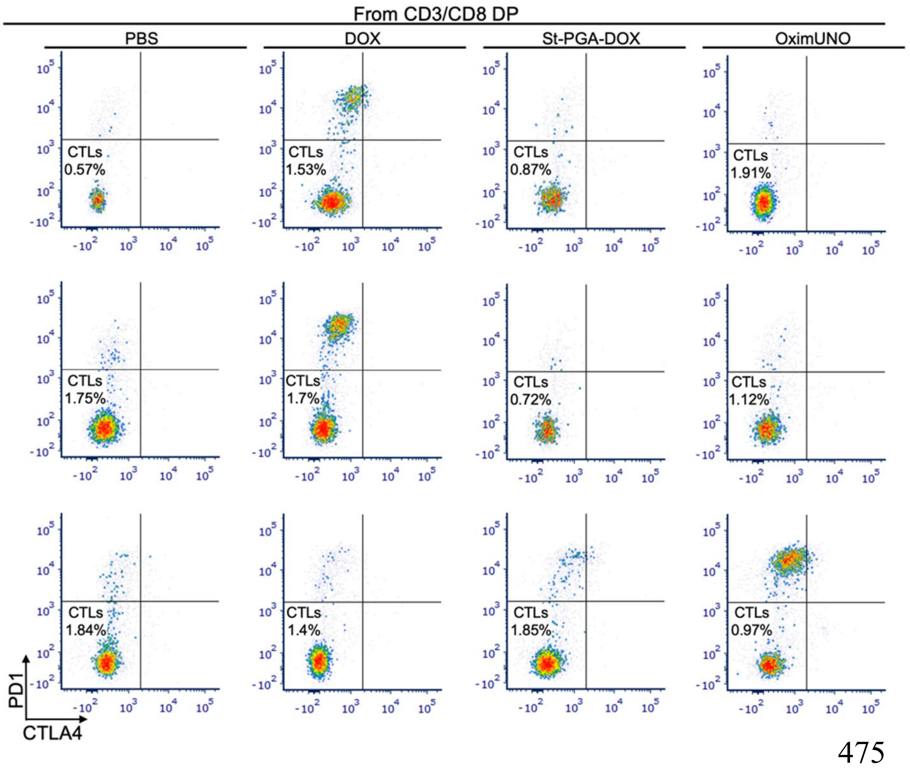


**Fig. S15. Flow cytometry gating for cytotoxic T lymphocytes (CTLs).** Cytometry plots for the CTLs populations of the treatment study shown in Figure 5. Percentages shown are from total cells. DP: double positive

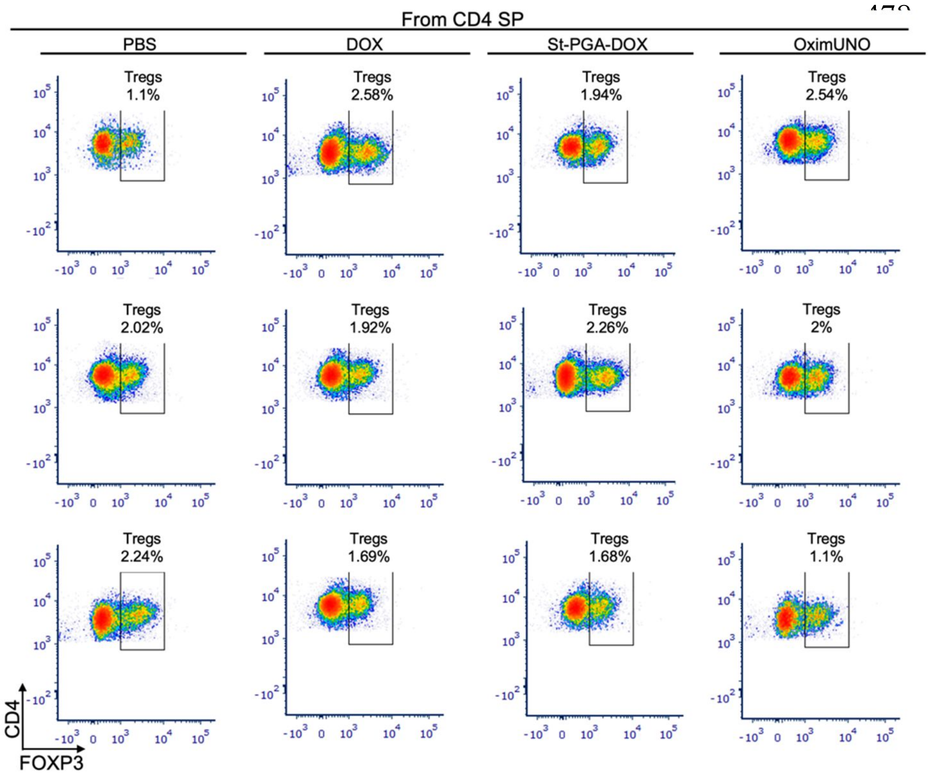


**Fig. S16. Flow cytometry gating for T regulatory cells (Tregs).** Cytometry plots for the Treg populations of the treatment study shown in Figure 5. Percentages shown are from total cells. SP: single positive

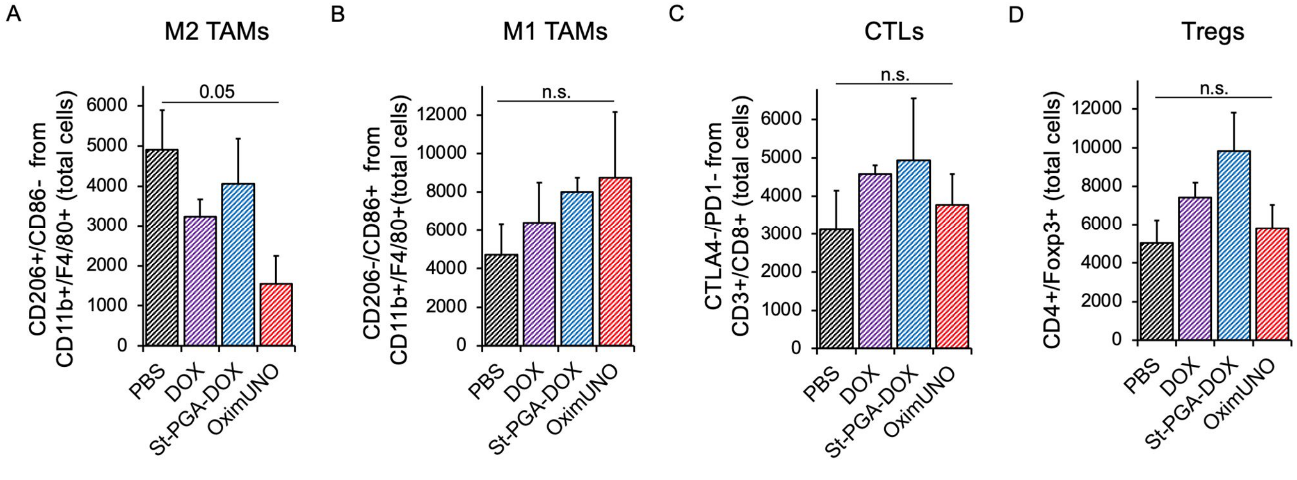


**Fig. S17. Flow cytometry analysis showing total cells.** Flow cytometry analysis showing total cells for (**A**) M2 TAMs, (**B**) M1 TAMs, (**C**) CTLs, and (**D**) Tregs. Error bars represent the SE of the mean.

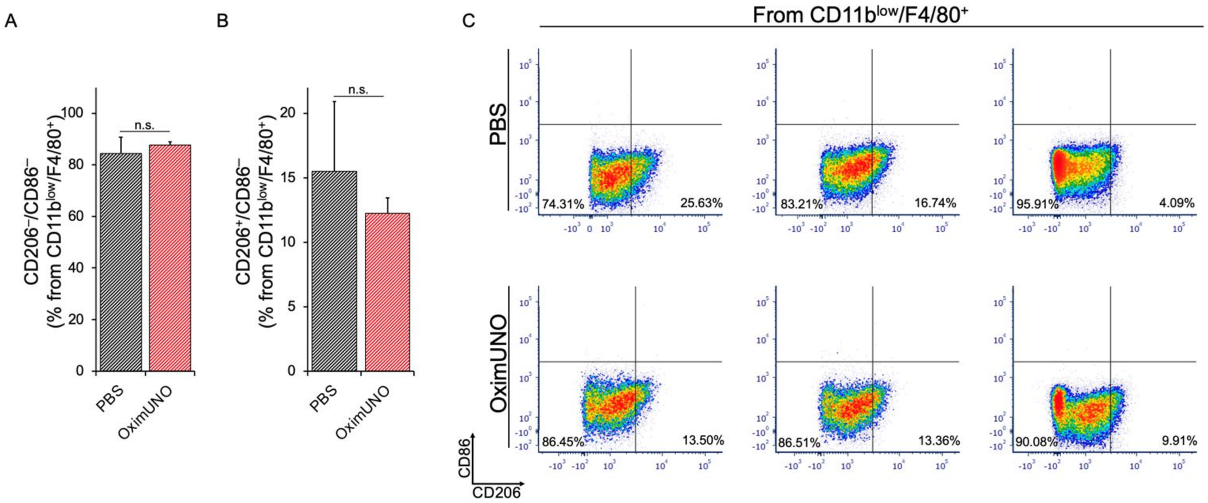


**Fig. S18. Treatment with OximUNO does not affect splenic macrophages. (A)** Percentages of $CD206^-/CD86^-$ splenic macrophages from $CD11b^{low}/F4/80^+$ cells, after treatment with OximUNO (from treatment study of Figure 5). **(B)** Percentages of $CD206^+/CD86^-$ splenic macrophages from $CD11b^{low}/F4/80^+$ cells after the treatment with OximUNO (from treatment study of Figure 5). **(C)** Flow cytometry plots for $CD11b^{low}/F4/80^+$ cells from spleens from treatment study of Figure 5.

### REFERENCES


1. Barz, M., Duro-Castano, A. & Vicent, M. J. A versatile post-polymerization modification method for polyglutamic acid: synthesis of orthogonal reactive polyglutamates and their use in "click chemistry". *Polym. Chem.* **4**, 2989–2994 (2013).
2. Van Lysebetten, D. *et al.* Lipid-Polyglutamate Nanoparticle Vaccine Platform. *ACS Appl. Mater. Interfaces* **13**, 6011–6022 (2021).
3. Arroyo-Crespo, J. J. *et al.* Tumor microenvironment-targeted poly-L-glutamic acid-based combination conjugate for enhanced triple negative breast cancer treatment. *Biomaterials* **186**, 8–21 (2018).
